# Space-time logic of liver gene expression at sublobular scale

**DOI:** 10.1101/2020.03.05.976571

**Authors:** Colas Droin, Jakob El Kholtei, Keren Bahar Halpern, Clémence Hurni, Milena Rozenberg, Sapir Muvkadi, Shalev Itzkovitz, Felix Naef

**Author notes:** Corresponding authors: Shalev Itzkovitz, Department of Molecular Cell Biology, Weizmann Institute of Science, Rehovot 7610001, Israel; : Felix Naef, Institute of Bioengineering, School of Life Sciences, Ecole Polytechnique Fédérale de Lausanne, CH-1015 Lausanne, Switzerland. These authors contributed equally to this work (Colas Droin, Jakob El Kholtei, Keren Bahar Halpern).

## Abstract

The mammalian liver performs key physiological functions for maintaining energy and metabolic homeostasis. Liver tissue is both spatially structured and temporally orchestrated. Hepatocytes operate in repeating anatomical units termed lobules and different lobule zones perform distinct functions. The liver is also subject to extensive temporal regulation, orchestrated by the interplay of the circadian clock, systemic signals and feeding rhythms. Liver zonation was previously analyzed as a static phenomenon and liver chronobiology at the tissue level. Here, we use single-cell RNA-seq to investigate the interplay between gene regulation in space and time. Categorizing mRNA expression profiles using mixed-effect models and smFISH validations, we find that many genes in the liver are both zonated and rhythmic, most of them showing multiplicative space-time effects. Such dually regulated genes cover key hepatic functions such as lipid, carbohydrate and amino acid metabolism, but also genes not previously associated with liver zonation such as chaperones. Our data also suggest that rhythmic and localized expression of Wnt targets could be explained by rhythmically expressed Wnt ligands from non-parenchymal cells near the central vein. Core circadian clock genes are expressed in a non-zonated manner, indicating that the liver clock is robust to zonation. Together, our comprehensive scRNA-seq analysis revealed how liver function is compartmentalized spatio-temporally at the sub-lobular scale.

## Introduction

The liver is a vital organ maintaining body physiology and energy homeostasis. The liver carries out a broad range of functions related to carbohydrate and lipid metabolism, detoxification, bile acid biosynthesis and transport, cholesterol processing, xenobiotics biotransformation, and carrier proteins secretion. Notably, the liver performs catabolic and anabolic processing of lipids and amino acids and produces the majority of plasma proteins^1^. Liver tissue is highly structured on the cellular scale, being heterogeneous in both cell-type composition and microenvironment^2^. In fact, liver tissue is made up of millions of repeating anatomical and functional subunits, called lobules, which in mice contain hepatocytes arranged in about 15 concentric layers with a diameter of about 0.5mm^3,4^. On the portal side of the lobule, blood from the portal vein and the hepatic arteriole enters small capillaries called sinusoids and flows to the central vein. This is accompanied with gradients in oxygen concentration, nutrients and signaling along the portal-central axis, with the latter notably involving the Wnt pathway^5,6^. Due to this polarization, hepatocytes in different layers perform separate functions. This is accompanied with gradients of gene expression along the portal-central axis, with some genes expressed more strongly near the central vein, and vice versa for portally expressed genes. This phenomenon is termed *liver zonation*^1,7^.

Recently, Halpern et al. combined single-cell RNA-sequencing (scRNA-seq) of dissociated hepatocytes and single-molecule RNA fluorescence in situ hybridization (smFISH) to reconstruct spatial mRNA expression profiles along the portal-central axis^8^. This analysis revealed an unexpected breadth of spatial heterogeneity, with ∼50% of genes showing spatially non-uniform patterns. Among them, functions related to ammonia clearance, carbohydrate catabolic and anabolic processes, xenobiotics detoxification, bile acid and cholesterol synthesis, fatty acid metabolism, targets of the Wnt and Ras pathways, and hypoxia-induced genes were strongly zonated.

In addition to its spatial heterogeneity, liver physiology is also highly temporally dynamic. Chronobiology studies showed that temporally gated physiological and metabolic programs in the liver result from the complex interplay between the endogenous circadian liver oscillator, rhythmic systemic signals, and feeding/fasting cycles^9,10,11^. An intact circadian clock has repeatedly been demonstrated as key for healthy metabolism, also in humans^12^. In addition, the hepatocyte clock has specifically been shown to play a major role in the physiological coordination of nutritional signals and cell-cell communication (including non-hepatocytic cells) controlling rhythmic metabolism^13^. Temporal compartmentalization can prevent two opposite and incompatible processes from simultaneously occurring, for example, glucose is stored as glycogen following a meal and is later released into the blood circulation during fasting period to maintain homeostasis in plasma glucose levels. Functional genomics studies of the circadian liver were typically performed on bulk liver tissue^14^. In particular, several studies showed how both the circadian clock and the feeding fasting cycles pervasively drive rhythms of gene expression in bulk, impacting key sectors of liver physiology such as glucose homeostasis, lipid and steroid metabolism^15–18^.

Here, we asked how these spatial and temporal regulatory programs interact on the level of individual genes and liver functions more generally. In particular, can zonated gene expression patterns be temporally modulated on a 24 h time scale? And conversely, can rhythmic gene expression patterns observed in bulk samples exhibit sub-lobular structure? More complex situations may also be envisaged, such as time-dependent zonation patterns of mRNA expression (or, equivalently, zone-dependent rhythmic patterns), or sublobular oscillations that would escape detection on the bulk level due to cancelations. On the physiological level, it is of interest to establish how hepatic functions might be compartmentalized both in space and time. To study both the spatial and temporal axes, we performed scRNA-seq of hepatocytes at four different times along the 24 h day, extending a previous approach^8,19^ to reconstruct spatial profiles at each time point. The resulting space-time patterns were statistically classified using a mixed-effect model describing both spatial and temporal variations in mRNA levels. In total, ∼5000 liver genes were classified based on their spatio-temporal expression profiles, and a few representative profiles were further analyzed with smFISH. Overall, this approach revealed the richness of space-time gene expression dynamics of the liver and provides a comprehensive view on how spatio-temporal compartmentalization is utilized at the sub-lobular scale in the mammalian liver.

## Results

### Single-cell RNA-seq captures spatiotemporal gene expression patterns in mouse liver

To investigate spatio-temporal gene expression patterns in mouse liver, we sequenced mRNA from individual liver cells obtained via perfusion from 10 *ad libitum* fed mice at 4 different times of the day (ZT = 0h, 6h, 12h and 18h, two to three replicates per time point). The interval between ZT0 and ZT12 has the light on and corresponds to the fasting period in mice, while feeding happens predominantly between ZT12 and ZT0. We here focused on hepatocytes by enrichment of cells according to size and *in silico* filtering, yielding a total of 19663 cells (Methods). To validate that the obtained scRNA-seq data captured the expected variability in both spatial and temporal mRNA levels, we generated a clustering analysis of all cells using a standard 2-dimensional t-SNE dimensionality reduction (Methods) and colored cells either by their positions along the central-portal axis (the a posteriori assigned layers, see below) (Figure 1A) or time (Figure 1B). The clustering revealed that portally and centrally expressed landmark transcripts, such as the cytochrome P450 oxygenases *Cyp2f2* and *Cyp2e1* involved in xenobiotics metabolism, mark cells in opposite regions of the projections (Figure 1C-D). Likewise, time-of-day gene expression varied along an orthogonal direction (Figure 1B), as shown for the fatty acid elongase *Elovl3* peaking at ZT0 (Figure 1E).

**Figure 1.**
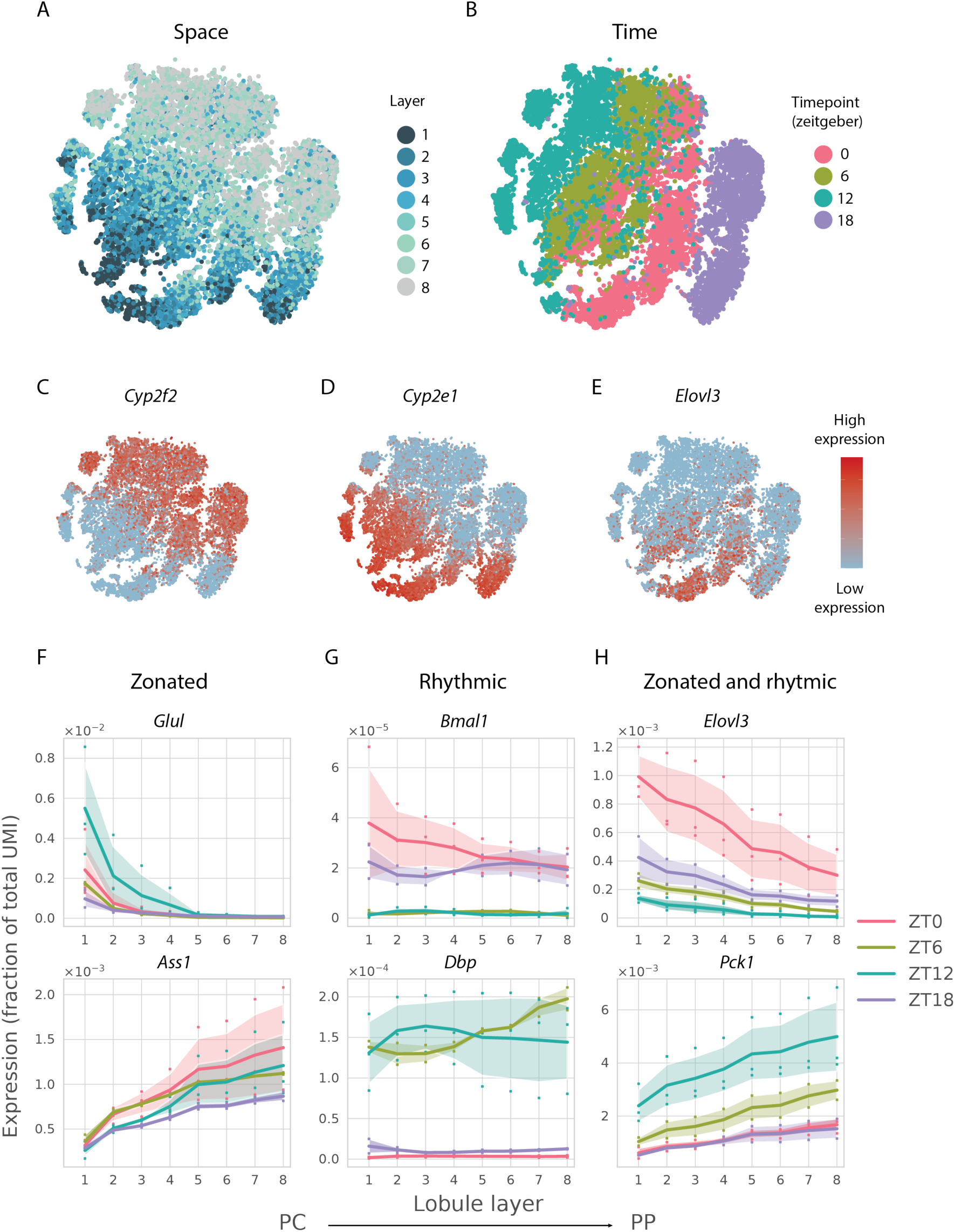
A scRNA-seq approach to space-time gene expression in mouse liver. **(A-E)** Global gene expression varies in both space and time, as shown using *t*-SNE visualizations of the scRNA-seq (19663 hepatocytes from 10 mice). Each dot represents one cell. Individual cells are colored by the (*a posteriori* assigned) lobule layer (A), Zeitgeber time (B), expression levels of the zonated genes *Cyp2f2* and *Cyp2e1* (C-D), or the temporally-regulated and centrally zonated gene *Elovl3* (E). **(F-H)** Reconstructed spatial profiles (lobule layers 1-8) of selected zonated genes (F, top: *Glul* pericentrally (PC) expressed, bottom: *Ass1* periportally (PP) expressed); rhythmic but non-zonated genes (G, top: core-clock gene *Bmal1* peaking at ZT18-0, bottom: clock-controlled *Dbp*, peaking at ZT6-12); zonated and rhythmic genes (H, Top: *Elovl3*, bottom: *Pck1*). Expression levels correspond to fraction of total UMI per cell in linear scale. Log-transformed profiles are in Supplementary Figure 1. Dots in F-H represent data points from the individual mice. Shaded areas represent one standard deviation (SD) across the mice.

To obtain spatial mRNA expression profiles for each gene along the central-portal axis, we here introduced eight lobule layers, to which we assigned each individual cell. For this, we adapted a previous method that uses expression levels of landmark zonated genes to define a central-portal coordinate^19^, with the modification that only landmark transcripts that were sufficiently expressed and that did not vary across mice and time points were used (27 central and 28 portal landmark genes, Methods). The resulting reconstructed (binned) mRNA expression profiles yielded 80 (8 layers over 10 mice) data points for each transcript. Although our resolution is lower compared with the typically 12-15 hepatocyte layers found in the liver^3,4^, these reconstructions faithfully captured reference zonated genes, with both central, and portal, expression (Figure 1F). Two examples of such genes are the centrally expressed glutamine synthethase (*Glul*), and the portally expressed urea cycle gene argininosuccinate synthetase (*Ass1*), showing mutually exclusive expression along the lobule^8^. The reconstruction also successfully identified transcripts of the core circadian clock, such as the master transcription factor *Bmal1* (also named *Arntl*), whose mRNA peaked between ZT18-ZT0 (Figure 1G)^20^. In addition to core clock genes, important clock outputs such as the PAR bZip transcription factor *Dbp*, which is a direct transcriptional target of BMAL1 regulating detoxification enzymes, peaked between ZT6-12 (Figure 1G)^21^. Finally, genes showing both zonated and rhythmic mRNA accumulation were found (Figure 1H), for example elongation of very long chain fatty acids 3 (*Elovl3*) is centrally expressed and peaks near ZT0, while phosphoenolpyruvate carboxykinase 1 (*Pck1*) regulating gluconeogenesis during fasting is expressed portally and peaks shortly before ZT12. Since most of the zonated profiles showed exponential shapes, and gene expression changes typically occur on a log scale^22^, we log-transformed the data for further analysis (Methods, Figure 1F and Supplementary Figure 1G-H-I). Together, these examples indicate that the obtained gene expression profiles reliably capture spatial and temporal regulation of hepatocyte gene expression.

### Space-time mRNA expression profiles categorized according to zonation and rhythmicity

To gain a systematic understanding of the space-time gene expression profiles, we next investigated if zonated gene expression patterns could be dynamic along the day, or conversely whether temporal expression patterns might be zone-dependent. To select a reliable set of reconstructed mRNA expression profiles for subsequent analyses, we filtered out lowly expressed genes as well as genes with significant biological variability across replicate liver samples, although this may be at the expense of a potentially decreased sensitivity (Methods). This yielded 5058 spatio-temporal gene expression profiles (Supplementary Figure 2A). An exploratory analysis of variance clearly identified zonated genes, rhythmic genes, and fewer genes showing variability along both axes, with known zonated and rhythmic genes distributed as expected (Figure 2A).

**Figure 2.**
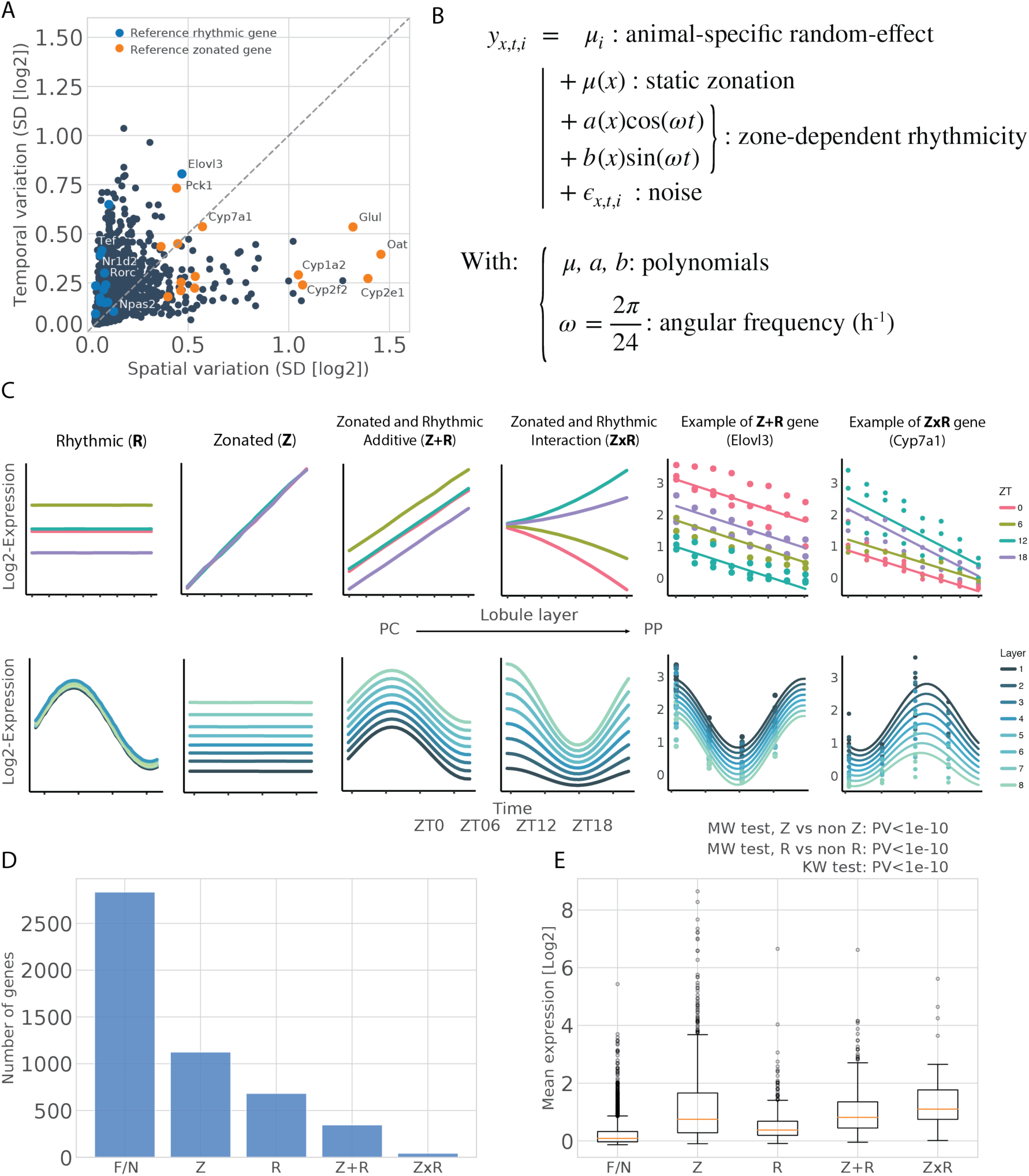
Space-time mRNA expression profiles categorized with mixed-effect models. **(A)** Spatial and temporal variations for mRNA transcript profiles, calculated as standard deviations (SD) of log2 expression along spatial or temporal dimensions. Colored dots correspond to reference zonated genes (orange) and reference rhythmic genes (blue) (Methods). **(B)** Extended harmonic regression model for spatio-temporal expression profiles describing a static but zonated layer-dependent mean *μ*(*x*), as well as layer-dependent harmonic coefficients (*a*(*x*)and *b*(*x*)). All layer-dependent coefficients are modeled as second order polynomials; *i*denotes the biological replicates. Temporal dependency is modelled with 24h-periodic harmonic functions. *μ*_*i*_ are random effects needed due to the data structure hierarchy (Methods). **(C)** Schema illustrating the different categories of profiles. Depending on which coefficients are non-zero (Methods), genes are assigned to: F/N (flat or noisy, not represented), Z (zonated), R (rhythmic), Z+R (additive zonation and rhythmicity), ZxR (interacting zonation and rhythmicity). Graphs emphasize either zonation (top), with the x-axis representing layers, or rhythmicity (bottom), with the x axis representing time (ZT). Right side of the panel: two examples of fits (*Elovl3* and *Cyp7a1*, respectively Z+R and ZxR). **(D)** Number of transcripts in each category. **(E)** Boxplot of the mean expression per category shows that zonated genes (Z, Z+R and ZxR) are more expressed than rhythmic (R) or flat/noisy (F/N). ZxR genes are the most expressed according to median expression (orange line). Box limits are lower and upper quartile, whiskers extend up to the first datum greater/lower than the upper/lower quartile plus 1.5 times the interquartile range. Remaining points are outliers. KW test stands for Kruskal-Wallis.

To identify possible dependencies between spatial and temporal variations, we built a mixed-effect linear model^23^ for the space-time mRNA profiles, which extends harmonic regression to include a spatial covariate (Figure 2B). In this model, rhythms are parameterized with cosine and sine functions, while spatial profiles are represented with (up to second order) polynomials. In its most complex form, the model uses nine parameters describing spatially modulated oscillations, and one intercept per mouse (Methods). When some of the parameters are zero, the model reduces to simpler mRNA profiles, for example purely spatial or purely temporal expression profiles (Figure 2C). We then used model selection^24^ to identify the optimal parameterization and category for each gene (Methods). *In fine*, we classified each mRNA profile into one of five types of patterns (Figure 2C). If only the intercept is used, the profile will be classified as flat or noisy (F/N, Methods). If only time-independent zonation parameters are retained, the predicted profile will be purely zonated (Z). If only layer-independent rhythmic parameters are retained, the predicted profile will be purely rhythmic (R). If only layer-independent rhythmic parameters and time-independent zonation parameters are retained, the profile is classified as independent rhythmic-zonated (Z+R). If at least one layer-dependent rhythmic parameter is selected, the profile will be termed interacting (ZxR). This classification revealed that, overall, about 30% of the mRNA profiles were zonated (Z, Z+R and ZxR) and about 20% were rhythmic (R, Z+R and ZxR) (Figure 2D). The peak times of these rhythmic transcripts were highly consistent with bulk chronobiology data^25^ (Supplementary Figure 2B). The entire analysis can be browsed as a web-app resource along with the corresponding data (https://czviz.epfl.ch).

Interestingly, we found that 7% of the analyzed genes in the liver were both zonated and rhythmic. Such dually regulated transcripts represent 25% of all zonated transcripts, and 36% of all rhythmic transcripts, respectively. For example, the previously shown *Elovl3* transcript, involved in fatty acid elongation, and *Pck1*, a rate limiting enzyme in gluconeogenesis, are prototypical Z+R genes (Figure 1H, Supplementary Figure 1H). Gluconeogenesis is an energetically-demanding task^3^. As mice are in a metabolically fasted state requiring glucose production towards the end of the light phase (∼ ZT10) and oxygen needed for ATP production is most abundant portally^26^, this process is indeed both spatially and temporally regulated. The dual regulation of zonated-rhythmic genes may therefore ensure optimal liver function under switching metabolic conditions.

Dually regulated genes were mostly Z+R, with only a minority of ZxR patterns. The average expression across categories showed that rhythmic genes are significantly less expressed on average than genes in zonated categories, likely reflecting shorter half-lives (Figure 2E and Supplementary Figure 2C). Surprisingly, we find few highly expressed flat genes. Together, we found that mRNA expression of many zonated genes in hepatocytes is not static, and is in fact compartmentalized both in space and time.

### Properties of dually zonated and rhythmic mRNA profiles

The majority of dually regulated genes are Z+R, which denotes additive (in log) space-time effects, or dynamic patterns where slopes or shapes of spatial patterns do not change with time (Figure 2C). On the other hand, interacting patterns (ZxR) are rare. Comparing the proportions of central, mid-lobular (peaking in the middle of the portal-central axis) and portal genes among the purely zonated genes (Z), and independently zonated and rhythmic genes (Z+R), did not reveal significant differences (Figure 3A), suggesting that rhythmicity is uncoupled with the direction of zonation. Similarly, comparing the phase distribution among the purely rhythmic genes (R) and the Z+R genes did not show a significant difference (Figure 3B), indicating that zonation does not bias peak expression time. Moreover, oscillatory amplitudes were uncorrelated with the zonation slopes in Z+R genes (Figure 3C). Finally, for ZxR genes with potentially more complex space-time patterns, we investigated the spreads in amplitudes and peak times across the layers (Figure 3D). For wave-like pattern (phase modulated profiles), the phase difference across the lobule was up to 3h, which corresponds to a difference in time between neighboring hepatocytes on the order of 10 minutes for lobules of about 15 cell layers. On the other hand, amplitude modulated patterns showed up to 2-fold difference in oscillatory amplitude across the lobule.

**Figure 3.**
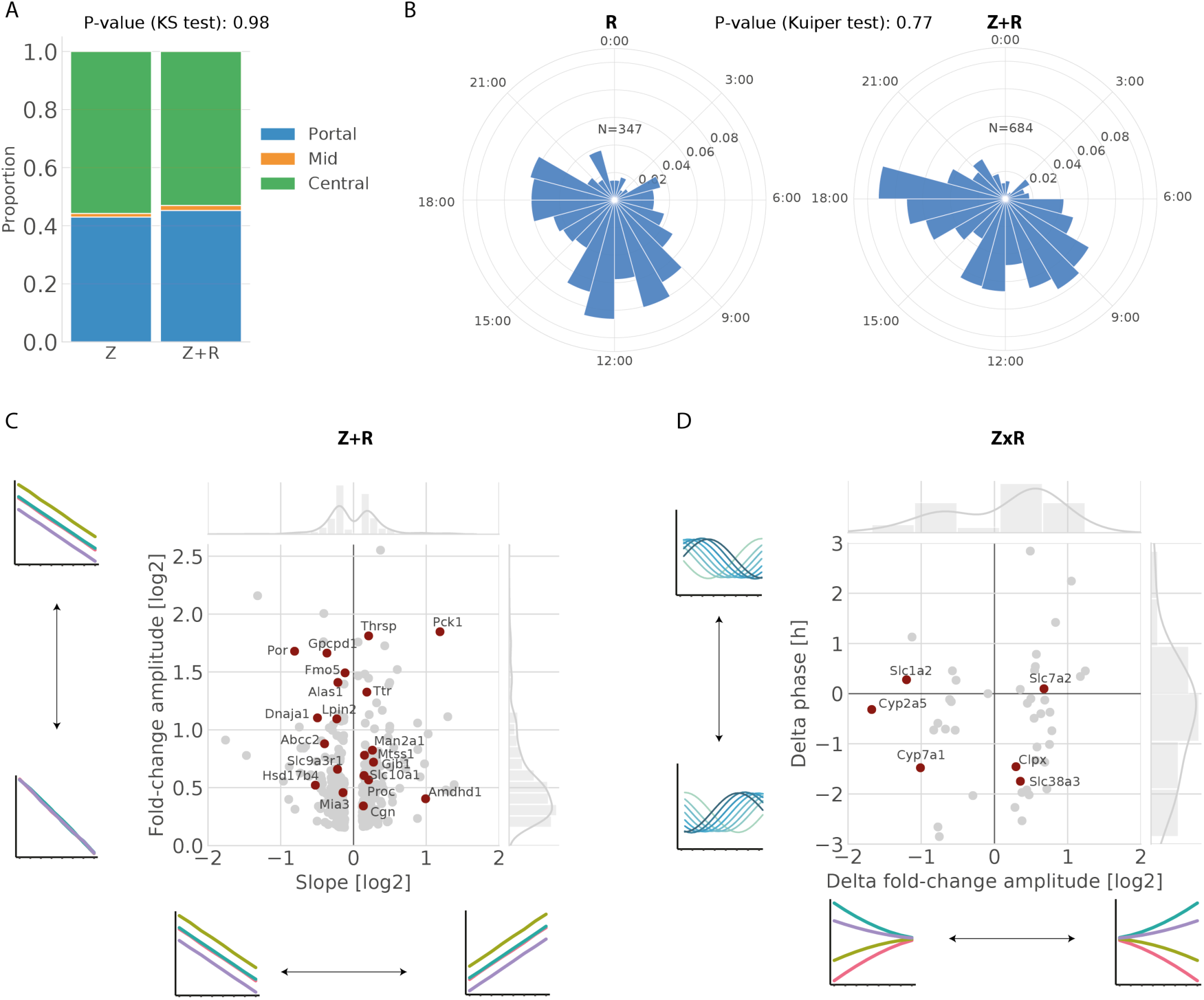
Properties of dually zonated and rhythmic mRNA profiles. **(A)** Proportions of pericentral (green) and periportal (blue) transcripts are similar in Z and Z+R. Mid-lobular genes (orange) are rare (<2%). **(B)** Peak time distributions of rhythmic transcripts are similar in R and Z+R categories. **(C-D)** Effect sizes of zonation (slope) *vs*. rhythmicity (fold-change amplitude, log2 peak-to-trough) in Z+R genes (C). Magnitude of time shifts (delta phase, in hours) *vs*. fold-change amplitude gradient (delta amplitude, in log2) along the central-portal axis in ZxR genes (D). Genes for which the protein is also rhythmic (p<0.05) in bulk data are indicated in dark red with the corresponding label (represented in Supplementary Figure 3).

To assess the potential physiological role of dually zonated and rhythmic transcripts, we asked if protein levels of the identified Z+R and ZxR genes accumulated rhythmically in a previous proteomics experiment^27^. In general, proteins rhythms are fewer, damped, and time-delayed compared to mRNA rhythms due to protein half-lives^14,27,28^ (Discussion). However, while R transcripts were twice more frequent than Z+R transcripts, the proportions were inverted for rhythmic proteins. Indeed, we found that among 65 rhythmic proteins (with q-value<0.2 in ref.^27^), 18 corresponded to Z+R and 10 to R transcripts. Moreover, the identified Z+R and ZxR genes with rhythmic protein accumulation cover key hepatic and zonated functions (Supplementary Figure 3, for a functional interpretation, see below) and include rate limiting enzymes. For example, for Z+R transcripts (Supplementary Figure 3A), PCK1 (rate limiting for gluconeogenesis), LPIN2 (*Lipin2*, catalyzes the conversion of phosphatidic acid to diacylglycerol during triglyceride, phosphatidylcholine and phosphatidylethanolamine biosynthesis), POR (cytochrome P450 oxidoreductase, required to activate P450 enzymes), DNAJA1 (HSP40 co-chaperone), ALAS1 (rate-limiting for heme biosynthesis), GNE (rate-limiting in the sialic acid biosynthetic pathway), THRSP (biosynthesis of triglycerides from medium-length fatty acid chains), show robust rhythms at the protein level. Similarly, for ZxR proteins, CYP7A1 (rate limiting enzyme in bile acid synthesis), CYP2A5 (coumarin 7-hydroxylase), SLC1A2 (high-affinity glutamate transporter), and multidrug resistance protein ABCC2 show rhythms on the protein level (Supplementary Figure 3B). Moreover, the protein rhythms accompanying those Z+R and ZxR transcripts peak with an expected delay of maximally about 6 hours^28^ compared to the mRNA peak times (Supplementary Figure 3C).

### smFISH analysis of space-time mRNA counts

To substantiate the RNA-seq profiles, we performed RNA single molecule fluorescence *in situ* hybridization (smFISH) experiments on a set of selected candidate genes with diverse spatio-temporal patterns. smFISH provides a sensitive and independent, albeit low-throughput, measurement of mRNA expression. Purely zonated genes (Z) were already well studied with smFISH^8^. To analyze the core-clock, we measured two genes peaking at different times, *Bmal1* and *Per1*, which were classified as R in the RNA-seq analysis. *Bmal1* (∼ZT0) and *Per1* (∼ZT12) phases were nearly identical in both experiments, and the rhythms did not depend on the lobular position consistent with R genes (Figure 4A). We analyzed three genes classified as Z+R: *Pck1* was indeed both portally biased and rhythmic in RNA-seq and smFISH (Figure 4B); *Elovl3* is both centrally biased and rhythmic in RNA-seq and smFISH, even though the amplitude of the oscillations was damped on the portal side in the FISH experiment (Supplementary Figure 4A); and for *Arg1 (Arginase 1*) the portal RNA-seq and smFISH profiles matched well (Supplementary Figure 4B). Finally, *Acly* showed a pattern in smFISH data which validates its classification as ZxR, with a lower amplitude on the portal side, where the transcript is more highly expressed (Figure 4C). Thus, overall, the reconstructed scRNA-seq and smFISH profiles were consistent, with minor discrepancies.

**Figure 4.**
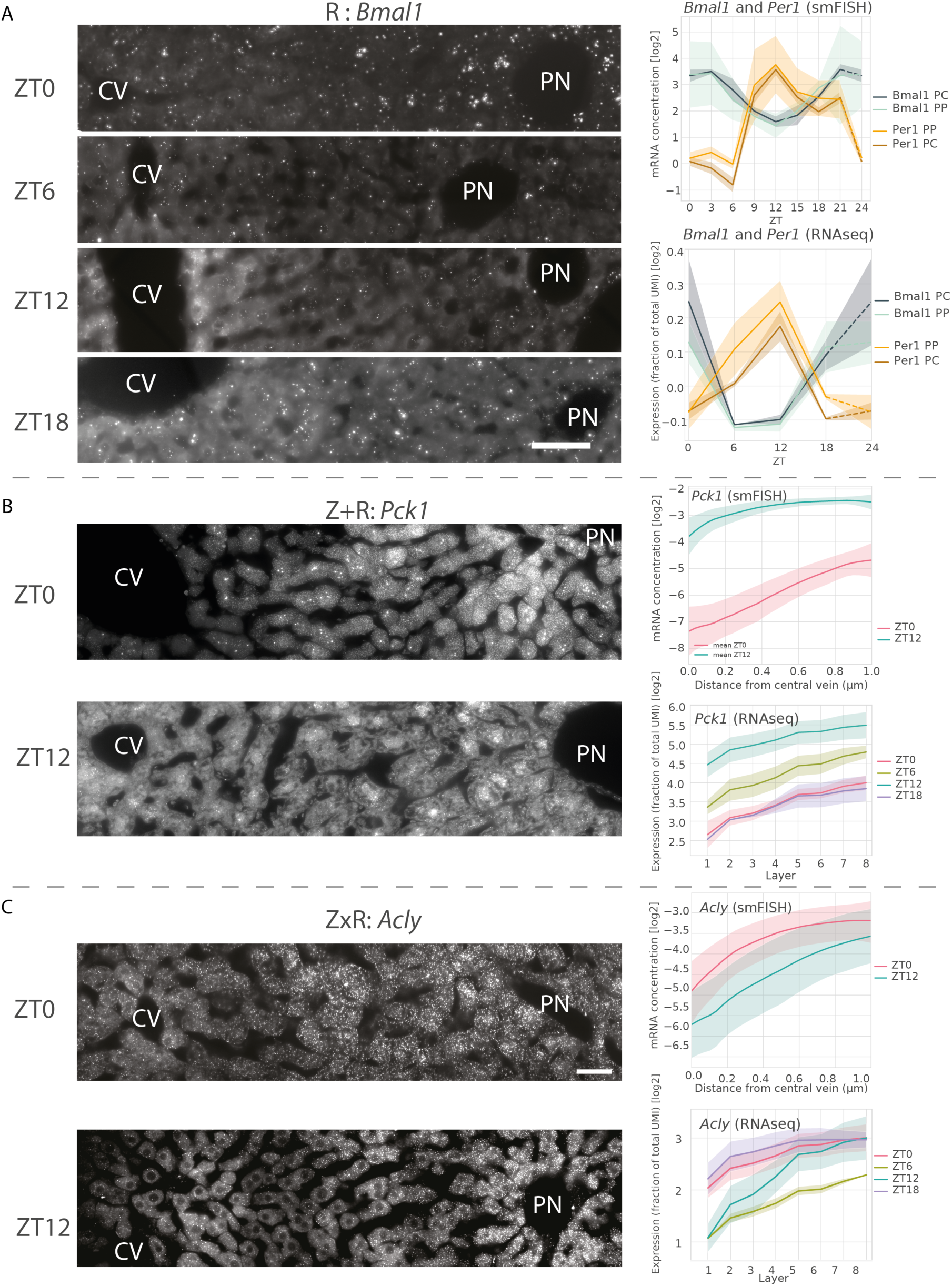
smFISH analysis of rhythmic and zonated transcripts. **(A)** smFISH (RNAscope, Methods) of the core clock genes *Bmal1* and *Per1* (both assigned to R) in liver slices sampled every 3 hours. **Left:** representative images at ZT0, ZT06, ZT12 and ZT18 for *Bmal1*. Central vein (CV) and a portal node (PN) are marked. Scale bar is 50µm. Endothelial cells lining the PC and cholangiocytes surrounding the PP were excluded from the quantification. mRNA transcripts and nuclei were detected in PN and PC zones (Methods). **Right:** temporal profiles of *Bmal1I* and *Per1* from smFISH (top, quantification of the number of mRNA transcripts at 8 time points ZT0 to ZT21, every 3 hours, shaded area indicate SD across images), in PN and PC regions, and scRNA-seq (bottom, shaded areas is SD across mice). **(B-C)** smFISH (Stellaris, Methods) for *Pck1* (Z+R) and *Acly* (ZxR). smFISH quantifications were made for ZT0 and ZT12 (Methods). **Left:** representative images at ZT0, ZT12 for *Pck1* (B) or *Acly* (C). Central vein (CV) and a portal node (PN) are marked. Scale bar - 20µm. **Right:** quantified profiles for each gene in the two time points from smFISH (top, shaded area indicate SD across images), and scRNA-seq data at the four time points (bottom, shaded areas is SD across mice).

### Space-time logic of hepatic functions

We next used our classification to explore the spatio-temporal dynamics of hepatic functions and signaling pathways in the liver. Given the prevalence of zonated gene expression profiles, we first analyzed if the circadian clock is sensitive to zonation. We found that profiles of reference core-clock genes (Supplementary Figure 5) were assigned to the rhythmic only category (R), except for *Cry1* and *Clock* that were assigned to Z+R, but with high probabilities also for R (Supplementary Table 2). This suggests that the circadian clock is largely non-zonated, as also seen in the smFISH (Figure 4A), and therefore robust to the heterogenous hepatic microenvironment.

**Figure 5.**
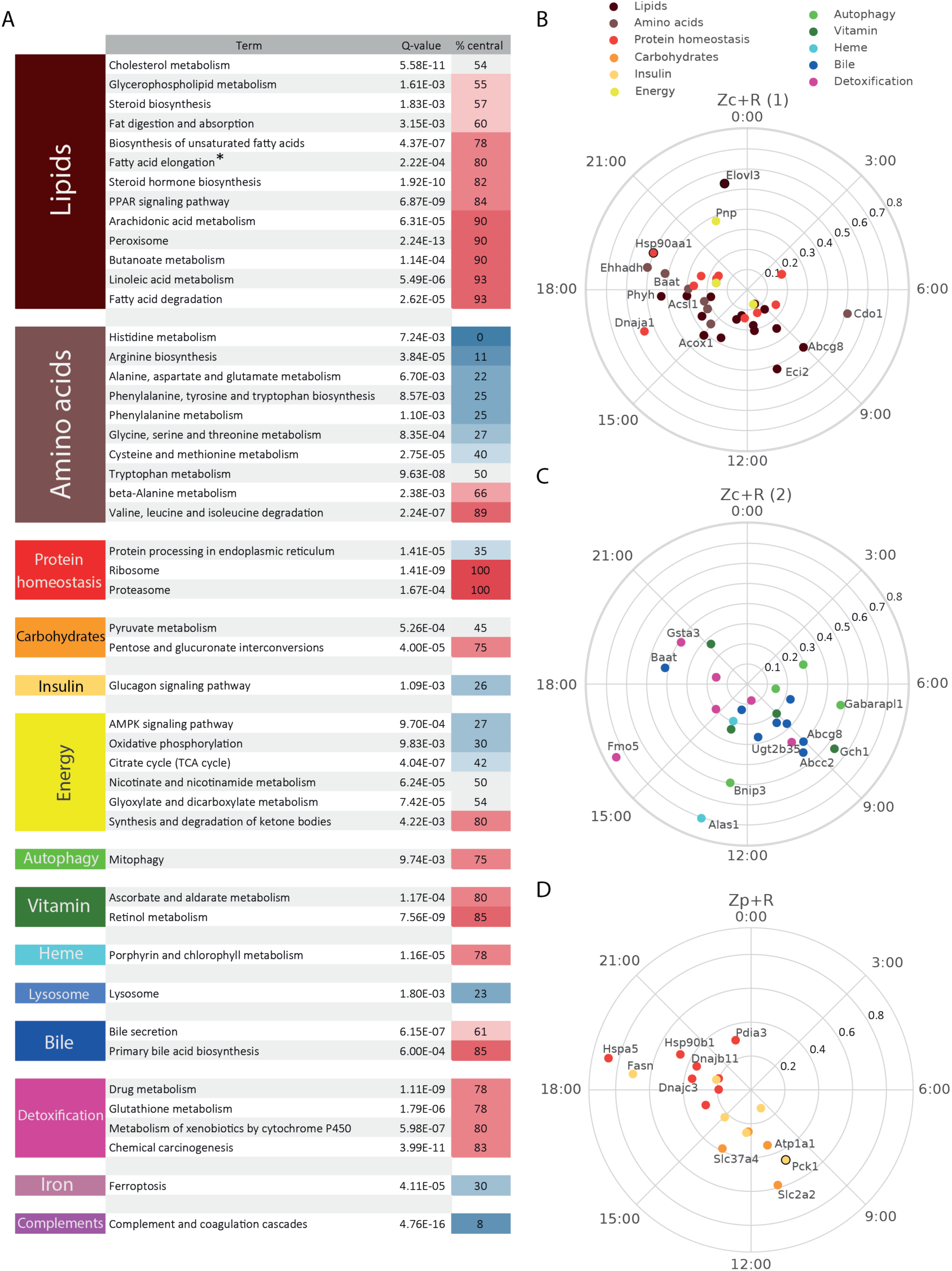
Space-time logic of compartmentalized hepatic functions for Z+R genes. **(A)** KEGG analysis of the Z and Z+R genes. For clarity, only KEGG pathways (second column) with multiple testing adjusted p-value<0.01 are presented (see Supplementary Table 3 all enriched functions). The percentage of central genes is represented by a blue-red gradient. *: appears central due to the KEGG annotation system; however, fatty acid elongation is biased portally (see text). **(B-C-D)** KEGG analysis of Z+R genes. Representations of genes in central (B-C) and portal (D) enriched Z+R categories (Supplementary Table 3). Polar representation: peak expression times are arranged clockwise (ZT0 on vertical position) and amplitudes (log2, values indicated on the radial axes) increase radially. The radius coordinate of genes with an amplitude >0.9 is halved (indicated with a black circle around the colored dot).

We then systematically explored enrichment of biological functions in the zonated category by querying the KEGG pathways database (Supplementary Table 3, Figure 5, Methods). In addition to recapitulating well documented zonated liver functions^8,29^, which we do not discuss here, this analysis highlighted Z and Z+R functions which, to our knowledge, had not been linked with liver zonation. For instance, we found that cytosolic chaperones accumulate centrally, while the endoplasmic reticulum (ER) chaperones linked with protein secretion accumulate portally (Figure 5B,D, Supplementary Figure 6). Both groups of chaperones peak during the activity/feeding phase, probably due to body temperature rhythms peaking during the active phase^30,31^, and likely reflect increased needs of protein folding during times of high protein synthesis. Also, we found that mRNAs of ribosomal protein genes accumulate centrally (Z), as do proteasome components (Figure 5A), the latter also containing rhythmic members (Z+R). In the liver, ribosomal proteins are rate-limiting for the synthesis of ribosomes, which themselves are rate-limiting for the synthesis of proteins^32^. Therefore, the overall protein synthesis rate is likely higher in these hepatocytes. Conversely, transcripts encoding components of the proteasome are involved in protein degradation. Together, these observations suggest that protein turnover is higher in centrally located hepatocytes, which are exposed to an environment with high concentrations of xenobiotics and hypoxic stress^33^.

**Figure 6.**
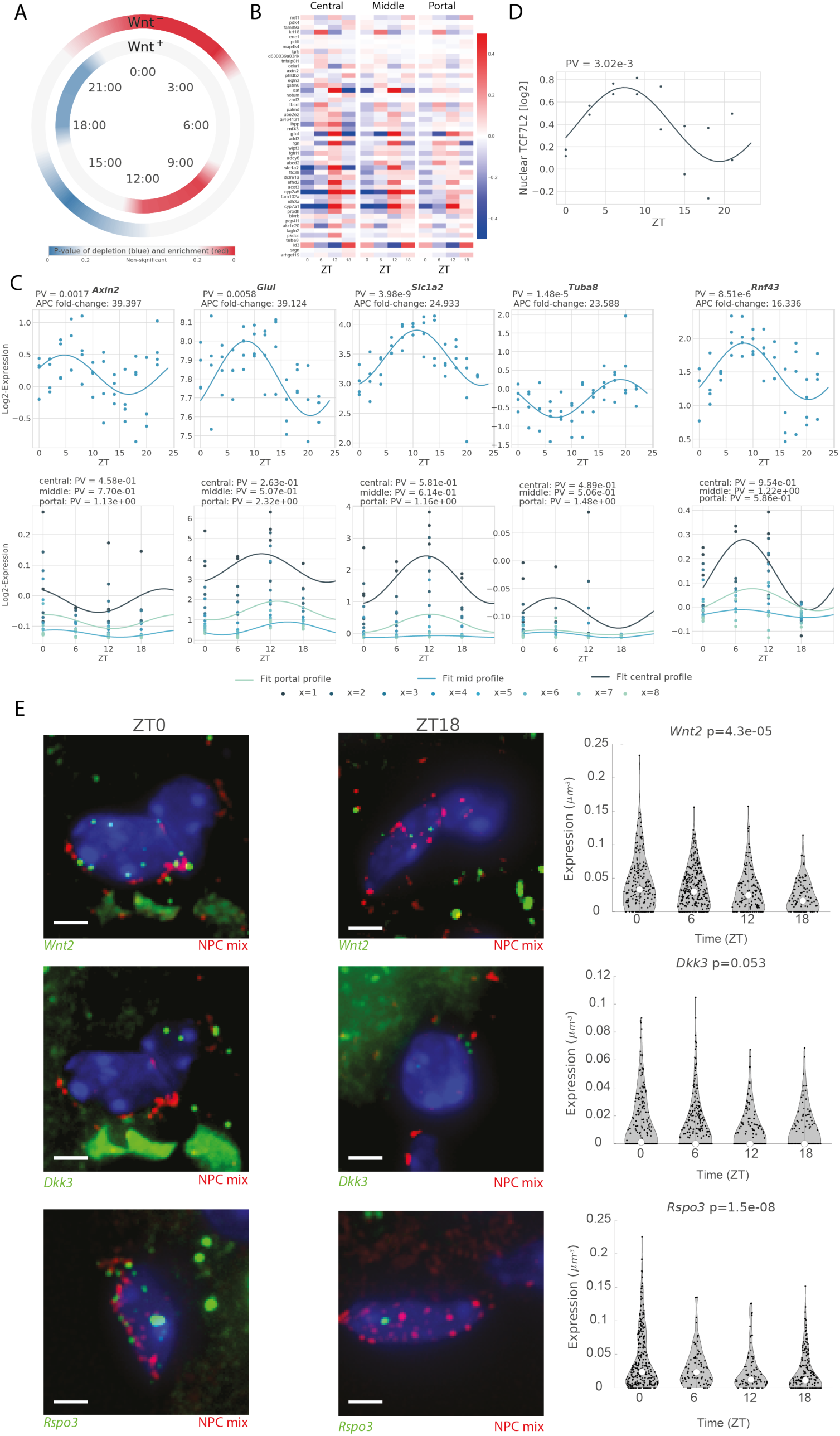
Rhythmic activity of Wnt signaling. **(A)** Enrichment/depletion at different times (window size: 3h), of both positive and negative Wnt targets (background: all R and Z+R genes). Colormap shows p-values (two-tailed hypergeometric test): red (blue) indicates enrichment (depletion). **(B)** Heatmaps representing scRNA-seq profiles of the top 50 Wnt targets (according to the Apc-KO fold change, *Lgr5* was also added) showing rhythmic mRNA in bulk (p<0.01, ref.^25^). The profiles are computed in three different zones of the central-portal axis: central (layers 1-2), mid-lobular (layers 3-4-5) and portal (layers 6-7-8). Gene profiles (log2) are mean-centered in each zone. An enrichment of the phases around ZT8-14 can be observed, in agreement with Figure 6A. **(C)** mRNA profiles from bulk RNA-seq (top) and scRNA-seq (bottom). Top five targets with the highest Apc-KO fold change, and rhythmicity in the bulk data (p<0.01, harmonic regression). Rhythmicity (indicated above each panel) is computed on bulk data or in three different zones for scRNA-seq data. **(D)** Nuclear protein abundance from ref.^35^ of the Wnt effector TCF4 (encoded by the *Tcf7l2* gene) in mouse liver shows a rhythm (p=0.003) peaking at ZT7.5, consistent with the accumulation of mRNA targets a few hours later (panel A). **(E) Left:** Representative smFISH images of *Wnt2, Dkk3*, and *Rspo3* expression at ZT0 (left) and ZT18 (right), shown in green. Markers of non-parenchymal cells (NPCs) are shown in red (Methods). Nuclei are stained in blue (DAPI). Scale bars, 2 µm; **Right:** Violin plots representing quantitative analysis of smFISH images (1420 cells of 189 central veins of at least two mice per time point). *Wnt2, Dkk3*, and *Rspo3* transcripts are quantified in NPCs lining the central vein (Methods). mRNA expression is in smFISH dots per µm^3^.

In addition, while many mitophagy genes are expressed centrally (Z), some of those also show robust temporal rhythms peaking during the fasting period (Z+R), in particular two gamma-aminobutyric acid receptor-associated proteins (*Gabarap* and *Gabarapl1*) with an important function in autophagosome mediated autophagy^34^. Consistent with this temporal regulation, we had previously reported that the nuclear abundance of the fasting-dependent regulators of autophagy TFEB and ZKSCAN3 peaked near ZT6^35^. Moreover, the centrally and synchronously accumulating Ubiquitin B mRNA (*Ubb*, Z+R) may contribute to triggering mitophagy^36^. Thus, centrally biased mitophagy may both participate in removal of damaged mitochondria in the stressed central environment.

Similarly, genes involved in bile acid synthesis and bile secretion are known to show zonated expression patterns^8^ (Figure 5C). Here, we found that while the rate-limiting enzyme in the bile acid biosynthetic pathway *Cyp7a1* (ZxR, Figure 2C) is known to be clock-controlled, with its mRNA^37,38^ and protein^39^ (Supplementary Figure 7D) expressed maximally early during the feeding period, the ABC transporters Sterolin-1 and 2 (ABCG5/ ABCG8, both identified as Z+R) which excrete most of the biliary cholesterol^40^ peak towards the end of the fasting period near ZT9.

Many detoxification enzymes of the cytochromes P450 (CYPs) superfamily are known to be centrally zonated in the liver^8^ and several of those are found in the Z+R category. In particular, the Flavin-containing monooxygenases Fmo1, 2, and 5, which are NADPH-dependent monooxygenases involved in drug and xenobiotics detoxification, exhibit Z+R mRNA patterns with peaks near ZT16 (Figure 5C). Also, the FMO5 protein accumulates rhythmically (Figure 3C, Supplementary Figure 3). Moreover, the rate limiting enzyme ALAS1 producing the P450 cofactor heme was found as a centrally zonated Z+R transcript with peak mRNA at ZT13, showing also a robust rhythm in protein expression (Supplementary Figure 3).

To substantiate the above finding on heat shock genes, we examined the space-time behavior of temperature-regulated genes. To this end, we considered targets (bound in ChIP-seq) of the heat shock transcription factor HSF1 (Chip-Atlas^41^, includes our own liver data^16^). This showed that several targets, known to peak during the active phase^30,31^, are also zonated (Supplementary Figure 6). Notably, the cytoplasmic *Hsp90aa1* (Hsp90A chaperone) and its interactor *Dnaja1* (HSP40), respectively mitochondrial *Hspd1* (HSP60), chaperones are expressed centrally where protein turnover is high. On the contrary, endoplasmic reticulum (ER) located chaperones: *Hspb1* (Hsp90B) and interactor *Pdia3*, as well as *Hspa5* (HSP70) and ER-resident *Dnajc3* and *Dnajb11* (DNAJ/HSP40) are expressed portally, consistent with their role in folding proteins in the secretory pathway (secretion is known to occur portally^29^). On the other hand, the analysis of cold induced genes (i.e. CIRBP and analogous, taken from ref.^42^) did not show zonated gene expression.

Finally, we note that among all KEEG pathway related to lipids, a majority show central enrichment (Figure 5A, Table S3). Inspection of the genes involved shows that this is due to the large number of genes related to peroxisomal β-oxidation, i.e. lipid catabolism, which are incidentally also listed in biosynthesis KEGG pathways (Table S3). However, fatty acid synthesis is biased portally, as supported by key portally expressed genes such as *<u>Fasn</u>, Srebf1, Acly, Acaca, Elovl2, Elovl5*, and which is consistent with the fact that oxygen needed for mitochondrial β-oxidation is most abundant portally^26^. On the other hand, *Elovl3*, which is known to be transcriptionally controlled by the peroxisome regulator PPARα and atypically regulated among Elov family fatty acid elongases^43^ is expressed centrally.

### Space-time logic of activity of signaling pathways

Signaling pathways that include Wnt, Ras and hypoxia have been shown to shape hepatocyte zonation^8^. We therefore examined the space-time activities of these pathways, extracted from the behavior of canonical target genes. We mainly focused on Wnt, as it is often considered the master regulator of liver zonation^44^. To systematically investigate spatio-temporal WNT/β-Catenin activity in liver, we extracted a set of Wnt targets derived from an APC KO in mouse liver^8,45^. We found that rhythmic transcripts (in the R, Z+R and ZxR categories) are enriched among targets of the Wnt pathway, showing a proportion that increases with the strength of the targets, with the strongest Wnt targets containing 80% of rhythmic transcripts (Supplementary Figure 7A). Positive Wnt targets were pericentral^8^ and peaked between ZT9 and ZT12, whereas negative Wnt targets were periportal^8^ and peaked between ZT21 and ZT3 (Figure 6A).

To obtain a temporal view of Wnt activity, we considered the top 50 Wnt pathway targets (according to the liver APC KO data) in the liver and analyzed the temporal profiles from high temporal resolution bulk liver mRNA and from our scRNA-seq binned in three different zones: central (layers 1-2), mid-lobular (layers 3-4-5) and portal (layers 6-7-8)(Figure 6B-C, Supplementary Figure 7B-C). This analysis confirmed that the peak times of rhythmic Wnt targets is preferentially between ZT9 and ZT12, and that the bulk and single cell data are consistent with each other, despite of the lower temporal sampling of the scRNA-seq. In fact, among the rhythmic genes detected in bulk RNA-seq, the five strongest Wnt targets were, in decreasing order: *Axin2, Glul, Slc1a2, Tuba8, Rnf43*, with the latter showing the largest amplitude (Figure 6C); all but *Tuba8* peaked in the morning. Note that *Glul*, an important marker of central zonation and canonical Wnt target (just like *Axin2* and *Rnf43*), was assigned to the Z category, but with second highest probability for ZxR (Supplementary Table 2), peaking at ZT12. Further evidence of rhythmic WNT/β-Catenin activity was provided by our previous proteomics data^35^ showing that the potent Wnt effector TCF4 (encoded by the *Tcf7l2* gene) has rhythmic nuclear abundance in mouse liver with a peak phase at ZT7.5 (Figure 6D), and hence explains the accumulation of its mRNA targets a few hours later.

In addition to the mRNA rhythms, we found that several of the Z+R or ZxR Wnt targets showed clear rhythms in bulk proteomics with the characteristic phase delays. The strongest five targets with rhythmic proteins (Supplementary Figure 7D) included the rate-limiting enzyme in the bile acid biosynthetic pathway CYP7A1, the NADPH-dependent monooxygenases involved in drug and xenobiotics detoxification FMO5, the P450 detoxification enzymes coumarin 7-hydroxylase (*Cyp2a5*), which may protect mice from dietary coumarin-induced toxicity^46^, the high-affinity glutamate transporter Slc1a2 (*Eaat2*), and the multidrug resistance protein ABCC2. All these proteins showed high amplitude protein rhythms peaking during the feeding phase between ZT12 and ZT18. Thus, Wnt transcription activity is clearly rhythmic in the liver, and this rhythm can propagate to protein expression.

We next asked whether the temporal oscillations in the expression of Wnt-activated genes might correlate with temporal oscillations in Wnt morphogens produced by pericentral non-parenchymal liver cells. To this end, we performed smFISH experiments and quantified the expression of the Wnt ligand *Wnt2*^5^ and of *Rspo3*^6,47^, a critical facilitator of Wnt signaling, as well as the Wnt antagonist *Dkk3*^19^ (Figure 6E). We found that both *Wnt2* and *Rspo3* in liver non-parenchymal cells (NPCs) exhibit non-uniform expression around the clock, with significantly higher mRNA levels at ZT0 (p=4×10^−5^ for *Wnt2*, and p=2×10^−8^ for *Rspo3*, Kruskal-Wallis, Figure 6E, right). Given various delays between mRNA accumulation of ligands and expression of the Wnt targets, this timing is compatible with the peak nuclear accumulation of the TCF4 (Tcf7l2 gene) transcription factor observed at ZT7.5 (Figure 6D) and with the peaks in Wnt-activated genes between ZT6-12 (Figure 6A-B). Differences in *Dkk3* expression were not significant (p=0.053). Thus, production of Wnt morphogens by central non-parenchymal liver cells might underlie the observed rhythmic Wnt pathway activity.

Ras signalling and hypoxia are two additional pathways that have been implicated in shaping hepatocyte zonation^8^. In agreement with ref.^8^, we found that the negative targets of Ras were enriched in central genes, whereas the positive Ras targets were enriched in portal genes (not shown). The rhythmic targets (R and Z+R) of hypoxia showed a pattern of temporal compartmentalization similar with those of Wnt (Supplementary Figure 7E): the negative targets were enriched around ZT0 (dark-light transition) and underrepresented around ZT14, while the positive targets were enriched around ZT10 and underrepresented around ZT3. Ras targets, positive or negative, did not exhibit significant temporal bias.

## Discussion

Recent genome-wide analyses of zonated gene expression in mouse and human liver^8,48,49^ uncovered a rich organization of liver functions in space at the sublobular scale, while chronobiology studies of bulk liver tissue revealed a complex landscape of rhythmic regulatory layers orchestrated by a circadian clock interacting with feeding-fasting cycles and systemic signals^35,50–52^. Here, we established how these two regulatory programs combine to shape the daily space-time dynamics of gene expression patterns and physiology in adult liver by extending our previous scRNA-seq approach^8^. We found that liver uses gene expression programs with many genes exhibiting compartmentalization in both space and time.

In this study, we chose to focus on the parenchymal cells in the liver, the hepatocytes, for which smFISH data on landmark zonated genes was readily available, which enabled reconstructing spatio-temporal mRNA profiles from scRNA-seq^8^. Zonation profiles of other cell types in the liver may be obtained as well; in fact, static zonation mRNA expression profiles have been obtained for liver endothelial cells, using a paired-cell approach^19^ in which mRNA from pairs of attached mouse cells were sequenced and gene expression from one cell type was used to infer the pairs’ tissue coordinates. In addition, *ab initio* reconstruction methods such as diffusion pseudo time^48^ or novoSpaRc^53^, in which a zonation coordinate is inferred by assuming that the major axis of variability for a cell type reflects transcriptome-wide gene expression changes associated with zonation, could be used for spatially sparse cell types with no available zonated marker genes, e.g. stellate or resident immune Kupffer cells. Moreover, it was recently found that rhythmic gene expression and metabolism in non-hepatocyte cells can be driven both by clocks in hepatocytes via cell-cell communication as well as feeding cycles^13^. Our computational framework for analyzing space-time logic of gene expression could be widely applicable in such future studies.

To study whether the observed space-time expression profiles may be regulated by either liver zonation, 24h rhythms in liver physiology, or both, we developed a mixed-effect model, combined with model selection. This enabled classifying gene profiles into five categories representing different modes of spatio-temporal regulation, from flat to wave-like. To validate these, we performed smFISH in intact liver tissue, which showed largely compatible profiles although some quantitative differences were observed. These differences most likely reflect the lower sensitivity of RNA-seq, uncertainties in the spatial analysis of smFISH in tissues, as well as known inter-animal variability in the physiologic states of individual livers, notably related to the animal-specific feeding patterns^25^.

Together, this temporal analysis confirms that a large proportion of gene expression in hepatocytes is zonated^8^ or rhythmic^17^, and in addition reveals marked spatio-temporal regulation of mRNA levels in mouse liver (Z+R and ZxR genes, comprising 7% of all detected genes according to our criteria). This means that zonated gene expression patterns can be temporally modulated on a circadian scale, or equivalently, that rhythmic gene expression profiles can exhibit sub-lobular structure. The dominant pattern for dually regulated gene was Z+R, which corresponds to additive effects of space and time in log, or multiplicate effects of gene expression levels, and describes genes expression profiles that are compartmentalized in both space and time. In other words, such patterns are characterized by shapes (in space) that remain invariant with time, but whose magnitudes are rhythmically rescaled in time. Or equivalently, the oscillatory amplitude (fold change) and phases are constant along the lobular coordinate, but the mean expression is patterned along the lobule. Such multiplicative effects could reflect the combined actions of transcriptional regulators for the zone and rhythm on promoters and enhancers of Z+R genes. Indeed, gene expression changes induced by several regulators combine multiplicatively^22^. Note that though the (relative) shape of Z+R patterns is invariant in time, threshold-dependent responses that would lie downstream of such genes would then acquire domain boundaries which can shift in time. Similar phenomena are expected for interacting profiles (phase and amplitude modulated) (ZxR) that we observed for a smaller number of genes.

As shown by us and others^27,28^, rhythms at the protein level are typically damped and phase-delayed compared the cognate mRNA rhythms, depending on the protein half-lives. Indeed, longer protein half-lives imply smaller oscillatory amplitudes and longer delays between mRNA and protein accumulation^14^. Analysis of liver proteomes in bulk showed that the number of cyclic proteins is significantly lower than the number of cyclic mRNAs, with delays approaching the predicted maximum of six hours. Here, we found genes from both Z+R and ZxR that exhibited rhythmic accumulation in bulk proteomics experiments, including genes encoding rate-limiting enzymes, suggesting that dually space-time regulated patterns have a physiological role in the liver. Moreover, phase delays between the mRNA and protein profiles were as expected. Future studies will utilize emerging spatial proteomics approaches to reconstruct a space-time liver proteomic atlas^54^.

In addition to previously discussed zonated liver functions^8^, a systematic querying of KEGG pathways highlighted Z+R functions not previously associated with rhythmic liver zonation. The roles and profiles of the corresponding genes allowed us to better understand the spatiotemporal logic of the identified pathways. For instance, we found that the expression levels of both ribosome protein genes (rate-limiting for protein synthesis^32^) and proteasome components (involved in protein degradation) were higher in central hepatocytes. Since the central environment is subject to high concentrations of xenobiotics and hypoxic stress, this could indicate an elevated protein turnover in this region, which would ensure that damaged proteins are rapidly exchanged with new, undamaged proteins. This interpretation is corroborated by the observed increased levels of cytosolic and ER chaperones during the feeding phase, to assist protein synthesis and secretion, thereby counteracting such protein stress.

It was previously shown that Wnt signaling can explain the zonation of up to a third of the zonated mRNAs^7^. Wnt ligands are secreted by pericentral non-parenchymal cells, mostly endothelial cells^5,6^, forming a graded spatial morphogenetic field. As a result, and as observed in our enrichment analysis, Wnt-activated genes were pericentrally-zonated. Moreover, both the scRNA-seq data and previous bulk mRNA and protein measurements showed that Wnt activity is rhythmic in the liver. Our smFISH analysis suggested that temporal fluctuations in the expression of *Wnt2* and *Rspo3*, two key Wnt ligands, secreted by pericentral non-parenchymal cells might underlie oscillatory and zonated expression of Wnt targets at times near the fasting/feeding transition.

In summary, we demonstrate how liver gene expression can be quantitatively investigated with spatial and temporal resolution and how liver functions are compartmentalized along these two axes. Our approach could be used to reconstruct spatio-temporal gene expression patterns in other zonated tissues such as the intestine and kidney^3^.

## Material and methods

### Animals and ethics statement

All animal care and handling were approved by the Institutional Animal Care and Use Committee of WIS and by the Canton de Vaud laws for animal protection (authorization VD3197.b). Male C57bl6 mice aged of 6 weeks, housed under reverse-phase cycle and under ad libitum feeding were used to generate sc-RNA-seq data of hepatocytes and single-molecule RNA-FISH (smFISH). Male mice between 8 to 10 weeks old, housed under 12:12 light-dark cycle, and having access to food only during the night (restricted-feeding) were used for smFISH of circadian clock genes.

### Hepatocytes isolation and single-cell RNA-seq

Liver cells were isolated using a modified version of the two-step collagenase perfusion method of Seglen^55^. The tissue was digested with Liberase Blendzyme 3 recombinant collagenase (Roche Diagnostics) according to the manufacturer instructions. To enrich for hepatocytes, we applied a centrifuge step at 30g for 3 min to pull down all hepatocytes while discarding most of the non-parenchymal cells that remained in the sup. We next enriched for live hepatocytes by 2 cycles of percoll gradient, hepatocytes pellet was resuspended in 25 ml of PBS, percoll was added for a final concentration of 45% and mixed with the hepatocytes. Dead cells were discarded after a centrifuge step (70g for 10min) cells were resuspended in 10x cells buffer (1x PBS, 0.04% BSA) and proceeded directly to the 10x pipeline. The cDNA library was prepared with the V2 chemistry of 10X genomics Chromium system according to manufactures instructions and sequencing was done with Illumina Nextseq 500 at estimated depth of 40,000 reads per cell.

Conceivably, the dissociation of liver tissue into individual cells and the purification of hepatocytes are relatively lengthy and may thus lead to alterations in mRNA expression. While it has been shown that mRNA levels do not change much during the purification and 24-hour cultivation of hepatocytes^56^, transcription rates on the other hand can be diminished by 8-fold to nearly 100-fold during this process^57^. This difference between nascent transcript and mature mRNA levels can be explained by the relatively long half-lives of liver-specific RNAs. In our case, the time needed from the dissociation of the tissue until the cell lysis is approximately 1 hour and the cell are not placed in culture. Since we are measuring mature transcripts, with half-lives in the range of typically 1-5 hours (Supplementary Figure 2C), the changes in mRNA levels due to the protocol will remain contained. In particular, the scRNA-seq of cells carry the typical hepatocyte gene expression signatures, for example, genes such as Alb or Apoa2 rank at 2nd and 5th position genome-wide. As further validation, we compared our reconstructed gene expression zonation profiles with the zonation profiles from the massively validated zonation study of ref.^8^, revealing a near-perfect agreement (Supplementary Figure 1F).

### Filtering of raw scRNA-seq data

The initial data analysis was done in R v3.4.2 using Seurat v2.1.0^58^. Each expression matrix was filtered separately to remove dead, dying and low-quality cells. We firstly only kept genes that were expressed in at least 5 cells in any of the ten samples. We then defined a set of valid cells with more than 500 expressed genes and between 1000 and 10000 unique molecular identifiers (UMIs) and secondly an additional expression matrix with cells having between 100 and 300 UMIs which was used for background estimation (Supplementary Figure 1A). Other UMI-filters have been tried, but yielded equal or less reliable profiles. The mean expression of each gene was then calculated for the background dataset and subtracted from the set of valid cells. This was subsequently filtered to only include hepatocytes by removing cells with expression of non-parenchymal liver cell genes. Next, the cells were filtered based on the fraction of mitochondrial gene expression. First, expression levels in each cell were normalized by the sum of all genes excluding mitochondrial and major urinary protein (*Mup*) genes. Indeed, as mitochondria are more abundant in periportal hepatocytes, the expression of mitochondrial genes is higher in this area^59^; and since these genes are very highly expressed, including them would reduce the relative expression of all other genes based on the cell’s lobular location. *Mup* genes are also highly abundant and mapping their reads to a reference sequence is unreliable due to their high sequence homology^60^. Moreover, *Mup* genes encode for pheromones that vary greatly between individuals to facilitate individual recognition^61^.

Mitochondrial content is often used to remove non-viable cells^62^. The mitochondrial content of sequenced hepatocytes exhibited a bi-modal behavior (Supplementary Figure 1B). To identify the range of mitochondrial fractions that included viable hepatocytes we used an intrinsic property of hepatocytes, which is the anti-correlation of the pericentral landmark gene *Cyp2e1* and the periportal landmark gene *Cyp2f2*^7^ (Supplementary Figure 1C). We found that hepatocytes with mitochondrial fraction in the range of 9-35% exhibited an almost perfect anticorrelation between *Cyp2e1* and *Cyp2f2* (Supplementary Figure 1D,E), suggesting that these are the best quality, and we consequently kept hepatocytes within this range of mitochondrial content for further analysis.

### t-SNE clustering

To validate that the expected spatial and temporal axes of variation are present in the scRNA-seq data, we generated a low-dimensional representation of all cells using the standard t-SNE (t-distributed stochastic neighbor embedding)^63^, a nonlinear dimensionality reduction technique that embeds high-dimensional data on a 2-dimensional plane such that points that are similar in high-dimensional space are close together on the 2-dimensional representation. We then colored cells either by their position along the central-portal axis, or by time of day.

### Spatial reconstruction of zonation profiles from scRNA-seq data

#### Choice of landmark genes

The reconstruction algorithm relies on *a priori* knowledge about the zonation of a small set of landmark genes to infer the location of the cells. Ref.^8^ used smFISH to determine the zonation pattern *in situ* of 6 such landmark genes and used them to reconstruct the spatial profiles of all other genes at a single time point. Since we here aimed at reconstructing zonation profiles at different time points, we could not rely on those landmark genes, which might be subject to temporal regulation. Therefore, we used an alternative strategy where we selected landmark zonated genes from ref.^8^ (q < 0.2), with the additional constraints that those should be highly expressed (mean expression in fraction UMI of more than 0.01% and less than 0.1%), and importantly vary little across mice and time. Specifically, we calculated the variability in the mean expression (across all layers) between all mice for every gene and removed genes with >= 10% variability. This yielded 27 central (*Akr1c6, Alad, Blvrb, C6, Car3, Ccdc107, Cml2, Cyp2c68, Cyp2d9, Cyp3a11, Entpd5, Fmo1, Gsta3, Gstm1, Gstm6, Gstt1, Hpd, Hsd17b10, Inmt, Iqgap2, Mgst1, Nrn1, Pex11a, Pon1, Psmd4, Slc22a1, Tex264*); and 28 portal (*Afm, Aldh1l1, Asl, Ass1, Atp5a1, Atp5g1, C8a, C8b, Ces3b, Cyp2f2, Elovl2, Fads1, Fbp1, Ftcd, Gm2a, Hpx, Hsd17b13, Ifitm3, Igf1, Igfals, Khk, Mug2, Pygl, Sepp1, Serpina1c, Serpina1e, Serpind1, Vtn*) landmark genes.

#### Reconstruction algorithm

The reconstruction algorithm is based on the algorithm in ref.^8^ and was used in the modified version from ref.^19^. The procedure was applied independently on each mouse, yielding ten spatial gene expression profiles for each gene, given as fraction of UMI per cell.

### Spatiotemporal analysis of liver gene expression profiles

#### Data

Each profile for the 14678 genes includes 8 layers from the pericentral to the periportal zone and 4 time points: ZT0 (n=3 biological replicates from individual mice), ZT6 (n=2), ZT12 (n=3) and ZT18 (n=2). The expression levels (noted as *x*) are then log-transformed as follows:

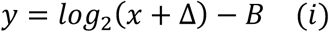

The offset Δ=10^−4^buffers variability in lowly expressed genes, while the shift *B*= −*log*_2_ (11×10^−5^)changes the scale so that *y* = 0 corresponds to about 10 mRNA copies per cell (we expect on the order of 1M mRNA transcripts per liver cell).

#### Reference genes

For ease of interpretation (Figure 2 and Supplementary Figure 2), we used a set of reference circadian genes and a set of reference zonated genes, highlighted in several figures.

The reference core circadian clock and clock output genes are the following: *Bmal1, Clock, Npas2, Nr1d1, Nr1d2, Per1, Per2, Cry1, Cry2, Dbp, Tef, Hlf, Elovl3, Rora, Rorc*.

The reference zonated genes are the following: *Glul, Ass1, Asl, Cyp2f2, Cyp1a2, Pck1, Cyp2e1, Cdh2, Cdh1, Cyp7a1, Acly, Alb, Oat, Aldob, Cps1*.

#### Gene expression variance in space and time

To analyze variability in space and time (Figure 2A) we computed, for each gene, the spatial variance *V*_*x*_ and the temporal variance *V*_*T*_. Let *y*_*x,t,j*_ represent the expression profile, with *j*the replicate index, *t* ∈ {1,2,…,*N*_*t*_}the time index, and *x* ∈ {1,2,…,*N*_*x*_}the layer index. Then, *V*_*x*_and *V*_*T*_ are computed as follows:

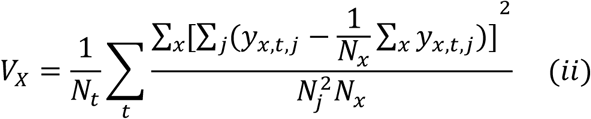

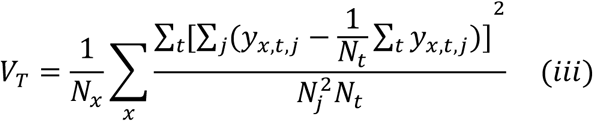

Thus, the spatial variance *V*_*x*_is computed along the space (and averaged over the replicates) for each time condition, and then averaged over time. The procedure is similar, symmetrically, for *V*_*t*_.

#### Gene filtering

For the analyses in Figure 2, we selected transcripts that were reproducible between replicates, as well as sufficiently highly expressed (see scatterplot in Supplementary Figure 2A). To assess reproducibility across replicates, we computed the average relative variance of the spatiotemporal profiles over the replicates:

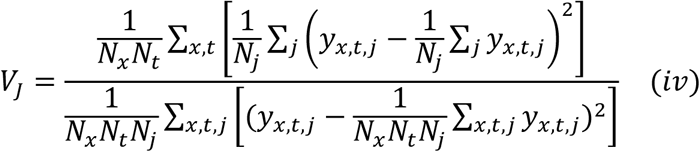

We considered genes with values below 50% (Supplementary Figure 2). To filter lowly expressed genes, we required the maximum expression level across layers and time points to exceed 10^−5^ (fraction of UMIs) which corresponds to y = 0 or about 10 copies of mRNA per cell. While this was quite more permissive than previous scRNA-seq studies it allowed to keep most reference circadian and zonated genes. However, scRNA-seq has still limited sensitivity and some potentially important genes may have been removed in the filtering process. In the end, our filters kept 5085 genes (1437 were removed due to low expression, 4733 due to high variance, and 3543 due to both), which were then used for subsequent analyses.

#### Mixed-effect model for spatiotemporal mRNA profiles

Since the data is longitudinal is space (8 layers measured in each animal), modelling the space-time profiles require the use of mixed-effect models. To systematically analyze the spatiotemporal mRNA profiles, we used a parameterized function. Specifically, the model uses sine and cosine functions for the time, and polynomials (up to degree 2) for space. Possible interaction between space and time are described as space-dependent oscillatory functions, or equivalently, time-dependent polynomial parameters. Our model for the transformed mRNA expression *y* reads:

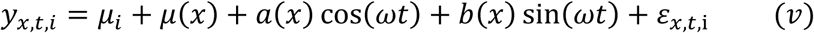

Here *t* is the time, *x* the spatial position along the liver layers, and *i*∈ {1,2,…,10}the animal index. This function naturally generalizes harmonic regression, often used for analysis of circadian gene expression^25^, by introducing space-dependent coefficients:

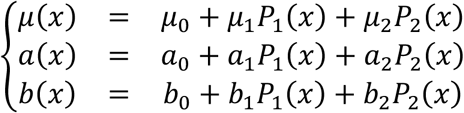

Here, *P*_1_and *P*_2_ are the Legendre polynomials of degrees 1 and 2, respectively; *μ*_0_, *μ*_1_and *μ*_2_ represent the static zonation profile, *a*_0_ and *b*_0_ represent the global (space-independent) rhythmicity of the gene, while *a*_1_, *a*_2_, *b*_1_, *b*_2_ represent layer-dependent rhythmicity. ε_*x,t,j*_ is a Gaussian noise term with standard deviation σ. In addition to the fixed-effect parameters described so far, we also introduced a mouse-specific random-effect *μ*_*i*_(with zero mean). This parameter groups the dependent layer measurements (obtained in the same animal) and thereby properly adjusts the biological sample size for the rhythmicity analysis.

Phases *φ*(related to peak times *t* through *t* = *φ*\*24/2*π*)and amplitudes *A*, for each profile can then be computed for any layer from the coefficients *a*(*x*)and *b*(*x*):

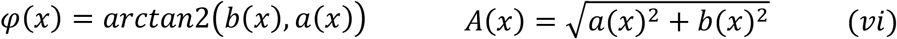

Note that peak-to-trough difference is 2*A*(*x*). The peak-to-trough ratio or fold change of the original expression levels is then 2^2*A*(*x*)^. We also note that an equivalent writing of the model formulates the problem in terms of time-dependent zonation parameters instead of space-dependent rhythmicity:

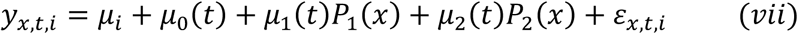

where:

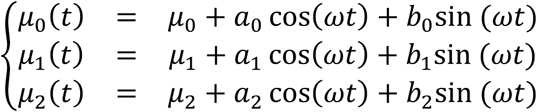

In this study, we fixed 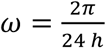 since the animals were entrained in a 24 h light-dark cycle and the low time resolution would prevent us from studying ultradian rhythms.

The model parameters, including the variance of the random effects and Gaussian noise strength σ, are estimated for each gene using the *fit* function from the Python library StatsModels (version 0.9.0). Nelder-Mead was chosen as the optimization method, and the use of a standard likelihood was favored over the REML likelihood to allow for model comparison^64^. To prevent overfitting of the gene profiles, we added a noise offset σ_0_= 0.15 [*log*2] to the estimated noise σ, in the expression of the likelihood function used in the mixed-effect model optimization.

Depending on the gene, the model presented in (*v*)and (*vii*)may be simplified by setting all or some of the (fixed) parameters to 0. For example, a non-oscillatory gene profile would normally have non-significant *a*_*j*_ and *b*_*j*_ parameters. In practice, considering the fixed effects, 2^9^ sub-models of various complexity can be generated. However, we added a few reasonable requirements to reduce the number of models. First, the intercept *μ*_0_ must be present in every model. Similarly, the parameters *a*_0_ and *b*_0_, providing a global rhythm, must be present in every rhythmic model. Finally, the parameters *a*_*j*_ and *b*_*j*_ for *j*=0,1,2 must be paired to ensure a proper phase definition (*vi*).

The models can then be classified in different categories, depending on the retained (non-zero) parameters (Figure 2C):

- The model comprising only the intercepts *μ*_0_ and *μ*_*i*_, termed flat or noisy (F/N).
- The models comprising only the intercepts and zonation parameters: *μ*_1_and/or *μ*_2_, termed purely zonated (Z).
- The models comprising only the intercepts and rhythmic parameters: *a*_0_ and *b*_0_, termed purely rhythmic (R).
- The models comprising only the intercepts, zonated parameters and rhythmic parameters: *μ*_1_and/or *μ*_2_, and *a*_0_, *b*_0_, termed independent (Z+R).
- The models comprising interaction parameters: *a*_*j*_ and *b*_*j*_ for *j*=1,2, termed interacting (ZxR).

Note that we only plot the fixed effects in the predicted gene profiles. The Bayesian Information Criterion (BIC) is then used for model selection, enabling to choose the most parsimonious model for each gene. Consequently, the F/N class also contains noisy profiles, since genes that are not well fitted with any complex model will then be assigned to the simplest model. Additionally, it appears that, for some profiles, several competing models can result in close BIC values (see e.g. the discussion on *Clock* and *Cry*1 in the Results). Therefore, when assigning hard classes, if some models have a relative difference of less than 1% in their BIC, we systematically keep the most complex model. Moreover, we also assigned probabilities to the different categories (F, Z, R, Z+R and ZxR), computed as Schwartz BIC weights^65^, which is useful in case of ambiguous classification (Supplementary Table 2). All best fits with their parameter values are listed in Supplementary Table 1.

#### Comparison of peak times with a bulk dataset

We compared the peak phases our rhythmically classified genes with those obtained from ref.^25^. These data consist of bulk liver RNA-seq data sampled every 2 hours for 24 hours, with 4 replicates per time condition. We only compared genes that for which rhythmicity is not changing across layers, *viz*. the R and Z+R categories. Note that since our dataset has a lower temporal resolution and fewer replicates per time point, we found overall less rhythmic genes.

To assess gene rhythmicity from ref.^25^, we used harmonic regression on the log-transformed profiles as previously. Using the same notation as above, we define the two following models:

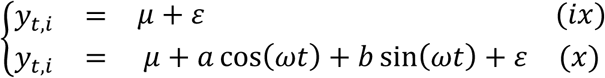

We then fit eq. (*viii*)and eq. (*ix*)to every transcript, and, for each of them, keep the model with the lowest BIC. We then compared the phases of transcripts classified as rhythmic in both datasets, and computed the circular correlation coefficient^66^.

### KEGG pathway Enrichment analysis

Functional annotation clustering from Enrichr^*67*^ for the categories F/N, Z and Z+R (which is then subdivided in central and portal), Zc+R (central zonated and rhythmic), Zp+R (portal zonated and rhythmic) and finally R was ran with standard parameters, using the standard KEGG 2019 Mouse set of pathways. The enriched pathways (adjusted p-value < 0.1) was then further annotated to compute, e.g. the number of central/portal genes in each category and the phase of each gene. This analysis is available Supplementary Table 3.

### smFISH

#### Analysis of Z+R and ZxR genes (Stellaris smFIH probes)

Preparation of probe libraries, hybridization procedure and imaging conditions were previously described^19^. smFISH probe libraries were coupled to TMR, Alexa594 or Cy5. Cell membranes were stained with alexa fluor 488 conjugated phalloidin (Rhenium A12379) that was added to GLOX buffer^68^. Portal node was identified morphologically on DAPI images based on bile ductile, central vein was identified using smFISH for Glul in TMR, included in all hybridizations. Images were taken as scans spanning the portal node to the central vein. Images were analyzed using ImageM^68^. Quantification of zonation profiles in different circadian time point were generated by counting dots and dividing the number of dots in radial layers spanning the portal-central axis by the layer volume.

#### Temporal analysis of circadian genes (RNA scope smFISH probes)

smFISH of R genes were done on fresh-frozen liver cryosections (8μm) embedded in O.C.T Compound (Tissue-Tek; Sakura-Finetek USA), sampled every three hours (ZT0 to ZT21). RNAscope® probes for *Bma1l* mRNA (Mm-Arntl, catalog #: 438748-C3) and *Per1* mRNA (Mm-Per1, catalog #: 438751) were used, according to the manufacturer’s instructions for the RNAscope Fluorescent Multiplex V1 Assay (Advanced Cell Diagnostics). To detect the central vein, an immunofluorescence of Glutamine Synthetase (ab49873, Abcam, diluted 1:2000 in PBS/BSA 0.5%/Triton-X0.01%) was done together with smFISH. Nuclei were counterstained with DAPI and sections were mounted with ProLong(tm) Gold Antifade Mountant. Liver sections were imaged with a Leica DM5500 widefield microscope and an oil-immersion x63 objective. Z-stacks were acquired (0.2μm between each Z position) and mRNA transcripts were quantified using ImageJ, as described previously in ref.^50^. Pericentral (PC) and Periportal (PP) veins were manually detected based on Glutamine Synthetase IF or on bile ducts (DAPI staining). The Euclidean distance between two veins and the distance from the vein of each mRNA transcript were calculated. mRNA transcripts were assigned to a PP or PC zone if the distance from the corresponding vein was smaller than one-third of the distance between the PP and PC veins (ranging from 50 to 130μm).

#### Wnt2, Rspo3 and Dkk3 expression in non-parenchymal cells (NPCs)

Preparation of probe libraries, hybridization protocol and imaging conditions were previously described^19^. The *Aqp1, Igfbp7 and Ptprb* probe libraries were coupled to TMR, the *Wnt2* library was coupled to *Alexa594* and the *Dkk3* or *Rspo3* library were coupled to *Cy5*. Cell membranes were stained with alexa fluor 488 coupled to phalloidin (Rhenium A12379) that was added to GLOX buffer^68^. The central vein was identified based on morphological features inspected in the DAPI and Phalloidin channels and presence of Wnt2-mRNA (detected by smFISH). Central vein niche NPCs were identified by co-staining of *Aqp1, Igfbp7* and *Ptprb*. The central vein area was imaged and the images were analyzed using ImageM^68^. We counted dots of *Wnt2, Rspo3* and *Dkk3* expression (corresponding to single mRNA molecules) in NPCs lining the central vein and removed background dots larger than 25 pixels. We then divided the dot count by the segmented cell volume. In total 1420 endothelial cells from 189 central veins of at least 2 mice per time point. In total 1420 NPCs from 189 central veins of at least 2 mice per time point (ZT0,6,12,18) were imaged and a Kruskal-Wallis test based on the mean mRNA dot concentration in each cell was performed to compare the ZT0 and ZT18 time points.

## Data availability

### scRNA-seq data

All scRNA-seq data is deposited in GEO with accession code GSE145197 (reviewer token xxx).

### Reconstructed gene profiles

Reconstructed spatio-temporal gene profiles are available as Matlab files at https://c4science.ch/diffusion/10261/

### Web-application

The whole dataset of gene profiles along with the analysis is available online as a web-application at the URL https://czviz.epfl.ch/. The application was built in Python using the library *Dash by Plotly* (version 1.0).

## Code availability

The code for fitting the mixed-effects models and generating the main figures is available at https://c4science.ch/diffusion/10261/

## Acknowledgments

This work was supported by the Rothschild Caesarea Foundation’ fund managed by Weizmann Institute and EPFL, a Swiss National Science Foundation Grant 310030_173079 (to F.N.), and the EPFL. S.I. is supported by the Henry Chanoch Krenter Institute for Biomedical Imaging and Genomics, The Leir Charitable Foundations, Richard Jakubskind Laboratory of Systems Biology, Cymerman-Jakubskind Prize, The Lord Sieff of Brimpton Memorial Fund, the Wolfson Foundation SCG, the Wolfson Family Charitable Trust, Edmond de Rothschild Foundations, the I-CORE program of the Planning and Budgeting Committee and the Israel Science Foundation (grants 1902/12 and 1796/12, the Israel Science Foundation grant no. 1486/16, the Chan Zuckerberg Initiative grant no. CZF2019-002434, the Broad Institute-Israel Science Foundation grant no. 2615/18, the European Research Council (ERC) under the European Union’s Horizon 2020 research and innovation program (grant agreement no. 768956, the Bert L. and N. Kuggie Vallee Foundation, and the Howard Hughes Medical Institute (HHMI) international research scholar award.

## Author contributions

F.N. and S.I. conceived the study. K.B.H., J.E.K. and C.H. prepared all the samples and performed the experiments. C.D. and F.N designed the modeling. K.B.H., J.E.K., C.H. and C.D. analyzed the data. M.R. and S.M. assisted with the scRNA-seq and smFISH experiments. F.N and S.I. supervised the study. C.D., J.E.K. and F.N. wrote the manuscript. All authors reviewed the manuscript and provided input.

## Ethics statement

All animal care and handling were approved by the Institutional Animal Care and Use Committee of Weizmann Institute of Science and by the Canton de Vaud laws for animal protection (authorization VD3197.b).

