## Supplementary Information for "Space-time logic of liver gene expression at sublobular scale"

#### Contents

**I Supplementary Figures** **2**

**II Supplementary Table captions** **10**

#### Part I
### Supplementary Figures

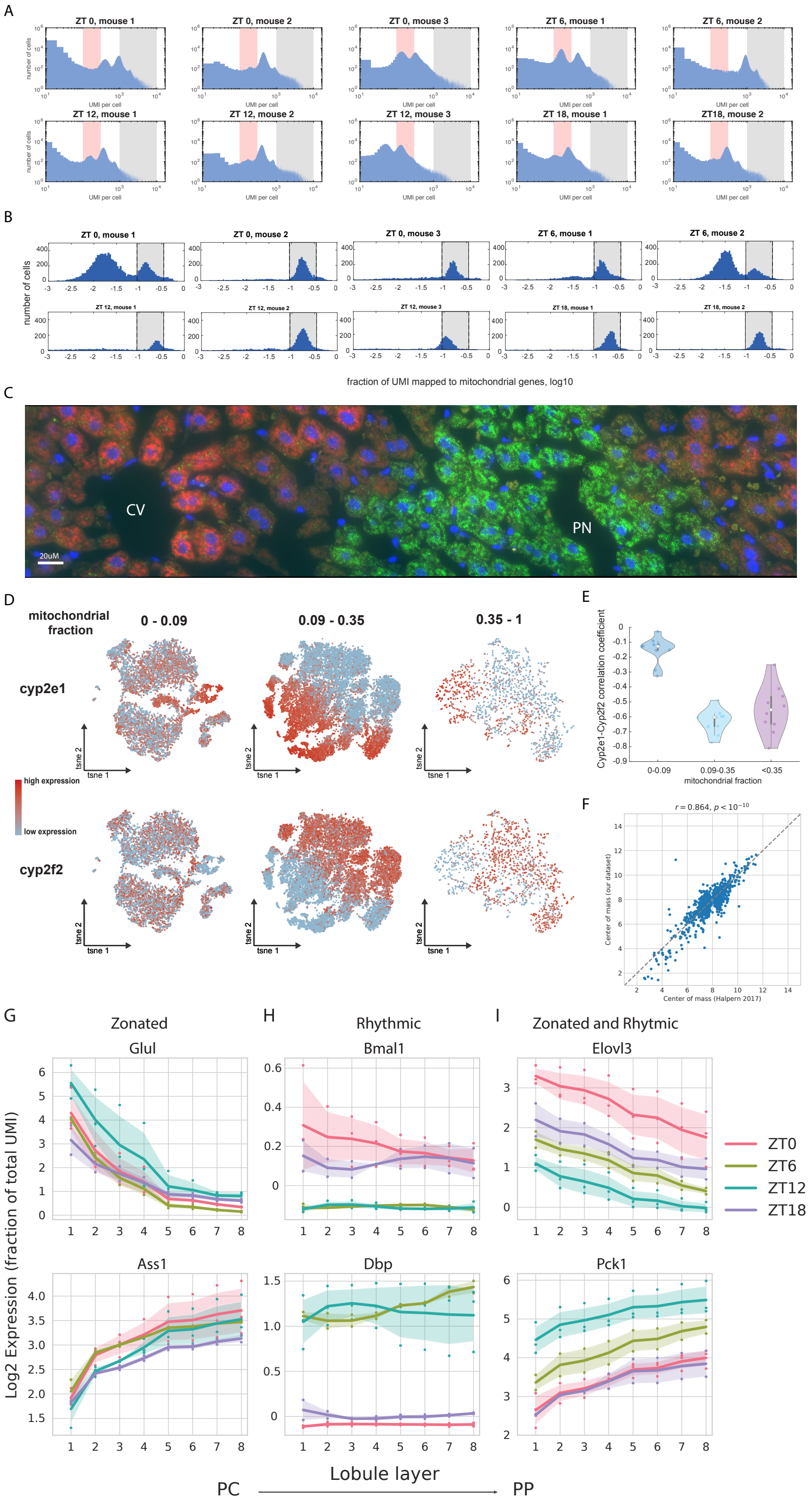
A
ZT 0, mouse 1
ZT 0, mouse 2
ZT 0, mouse 3
ZT 6, mouse 1
ZT 6, mouse 2
ZT 12, mouse 1
ZT 12, mouse 2
ZT 12, mouse 3
ZT 18, mouse 1
ZT18, mouse 2
number of cells
UMI per cell
B
number of cells
fraction of UMI mapped to mitochondrial genes, log10
C
CV
PN
20uM
D
mitochondrial fraction
0 - 0.09
0.09 - 0.35
0.35 - 1
cyp2e1
cyp2f2
tsne 1
tsne 2
high expression
low expression
E
Cyp2e1-Cyp2f2 correlation coefficient
0-0.09
0.09-0.35
<0.35
mitochondrial fraction
F
Center of mass (our dataset)
Center of mass (Halpern 2017)
G
Zonated
Glul
Ass1
H
Rhythmic
Bmal1
Dbp
I
Zonated and Rhytmic
Elovl3
Pck1
ZT0
ZT6
ZT12
ZT18
Log2 Expression (fraction of total UMI)
Lobule layer
PC
PP

**Supplementary Figure 1: scRNA-seq pre-processing.** (**A**) Histogram of number of UMIs per cell barcode for each mouse. Red patches mark the cells used for background estimation (100-300 UMI/cell barcode), gray patches mark the cells used for downstream analysis (1000-10000 UMI/cell barcode). (**B**) Histogram of fraction of all UMIs mapping to mitochondrial genes. Filter used for downstream analysis in grey (0.09-0.35). (**C**) smFISH staining of a representative liver lobule with probes against *Cyp2e1* (red) and *Cyp2f2* (green). CV = central vein, PN = portal node. (**D**) Expression of *Cyp2e1* and *Cyp2f2* in cells with different fraction of mitochondrial expression. Three different filters for the fraction of UMIs mapping to mitochondrial genes (0-0.09, 0.09-0.35, 0.35-1) were applied, the data of all mice merged and the resulting datasets visualized as t-SNE plots. (**E**) Violin plots for the correlations between *Cyp2e1* and *Cyp2f2* expression in single hepatocyte populations with different filters for fractions of mitochondrial expression. Each dot represents one mouse. (**F**) Comparison of the zonation profiles of Z and Z+R genes obtained in our current study and the previous reconstruction *Halpern et al. 2017.* Profiles were interpolated to fit 15 layers, where 1 is pericentral and 8 is periportal. Dots indicate the center of mass (expression-weighted lobule layer) of the Z and Z+R genes computed in both datasets, for gene having an average expression of at least $10^{-5}$ in *Halpern et al.*. $r$ is the correlation coefficient, $p$ the corresponding p-value. (**G-I**) Expression levels of the reconstructed profiles for the genes from Figure 1F-H after log-transformation (Methods). Shaded areas represent SD across mice.

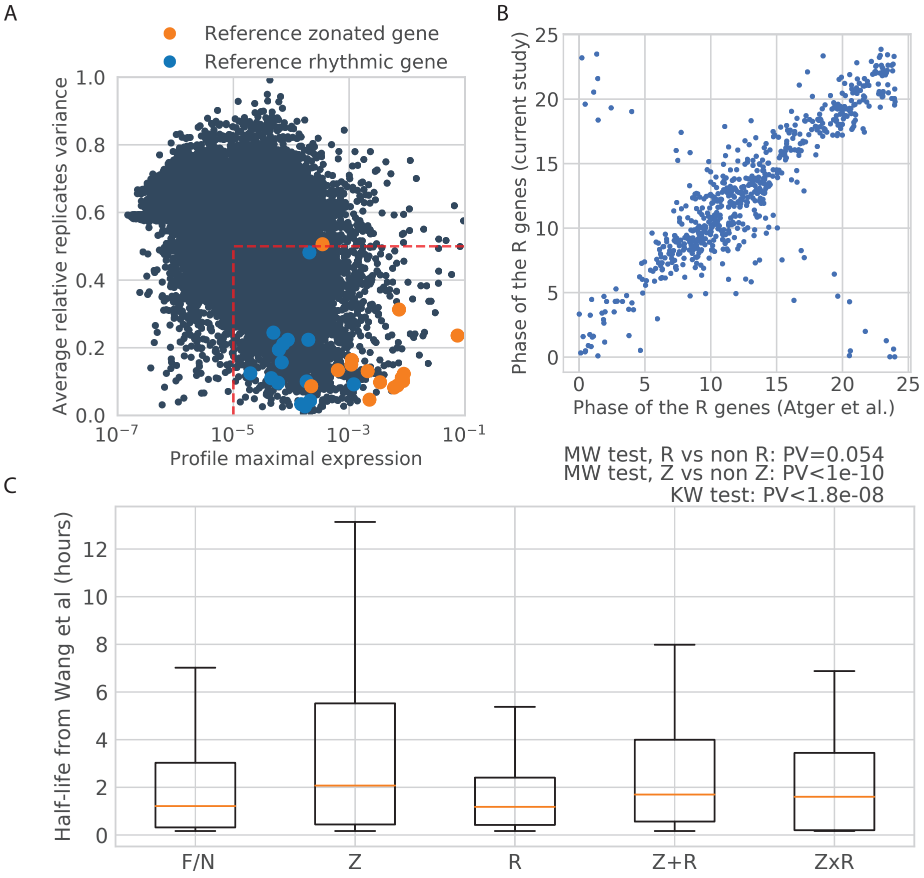


**Supplementary Figure 2: Pre-filtering of the genes and comparison with external datasets.** (**A**) Biological variability of gene profiles across independent replicate liver samples, quantified in terms of the average relative replicate variance. 0 shows perfectly reproducible profiles while 1 the most variable genes (Methods). Genes inside the bottom-right box (x-cutoff at $10^{-5}$; y-cutoff at 0.5) are selected and contain all but one of the reference genes. Colored dots show reference zonated genes (blue) and reference rhythmic genes (orange). (**B**) Comparison of the peak times for rhythmic genes in R and Z+R, with the dataset from *Atger et al, 2015*. Circular correlation coefficient is 0.746 (Methods). (**C**) Boxplot of the mRNA half-lives (data from *Wang, J. et al, 2017*) shows that R genes as a group (median, orange line) are the shortest-lived. Box limits are lower and upper quartiles, whiskers extend up to the first datum greater/lower than the upper/lower quartile plus 1.5 times the interquartile range. Remaining points are marked. MW stands for Mann-Whitney, KW stands for Kruskal-Wallis.

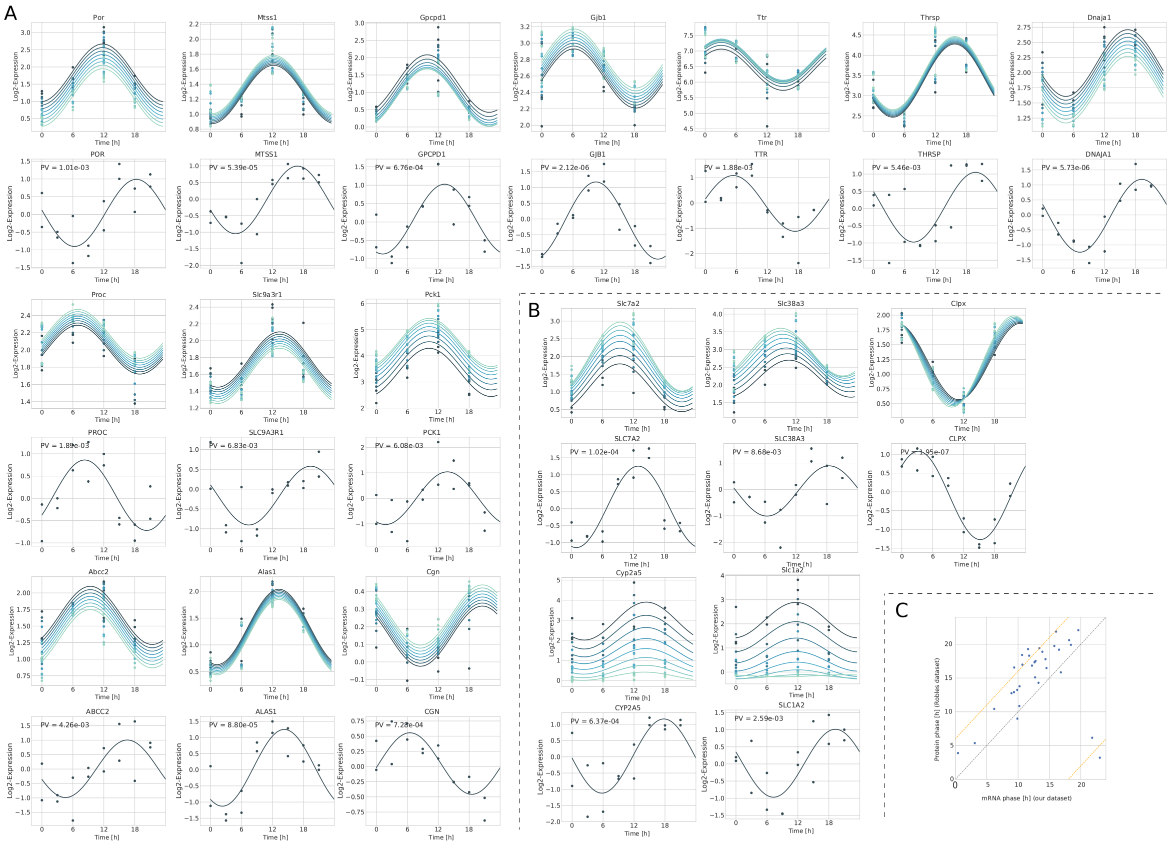


**Supplementary Figure 3: Z+R and ZxR transcripts with corresponding rhythmic protein accumulation in bulk mass spectrometry data.** (**A-B**) Rhythmic proteins corresponding to Z+R (A) and ZxR (B) transcripts were selected from *Robles et al., 2014*, from original Table S2), and fitted with a harmonic function (p-value indicated above the plot). Only proteins having a p-value¡0.01 are represented. (**C**) Scatter plot of the phase of the fits from the transcripts (x-axis) against the phase of the fits from the proteins (y-axis). The diagonal is indicated with a dashed grey line, the theoretical upper bound (6h) for the delay between mRNA and protein is indicated with a dashed red line. All rhythmic proteins (q¡0.2 in the original analysis) are represented.

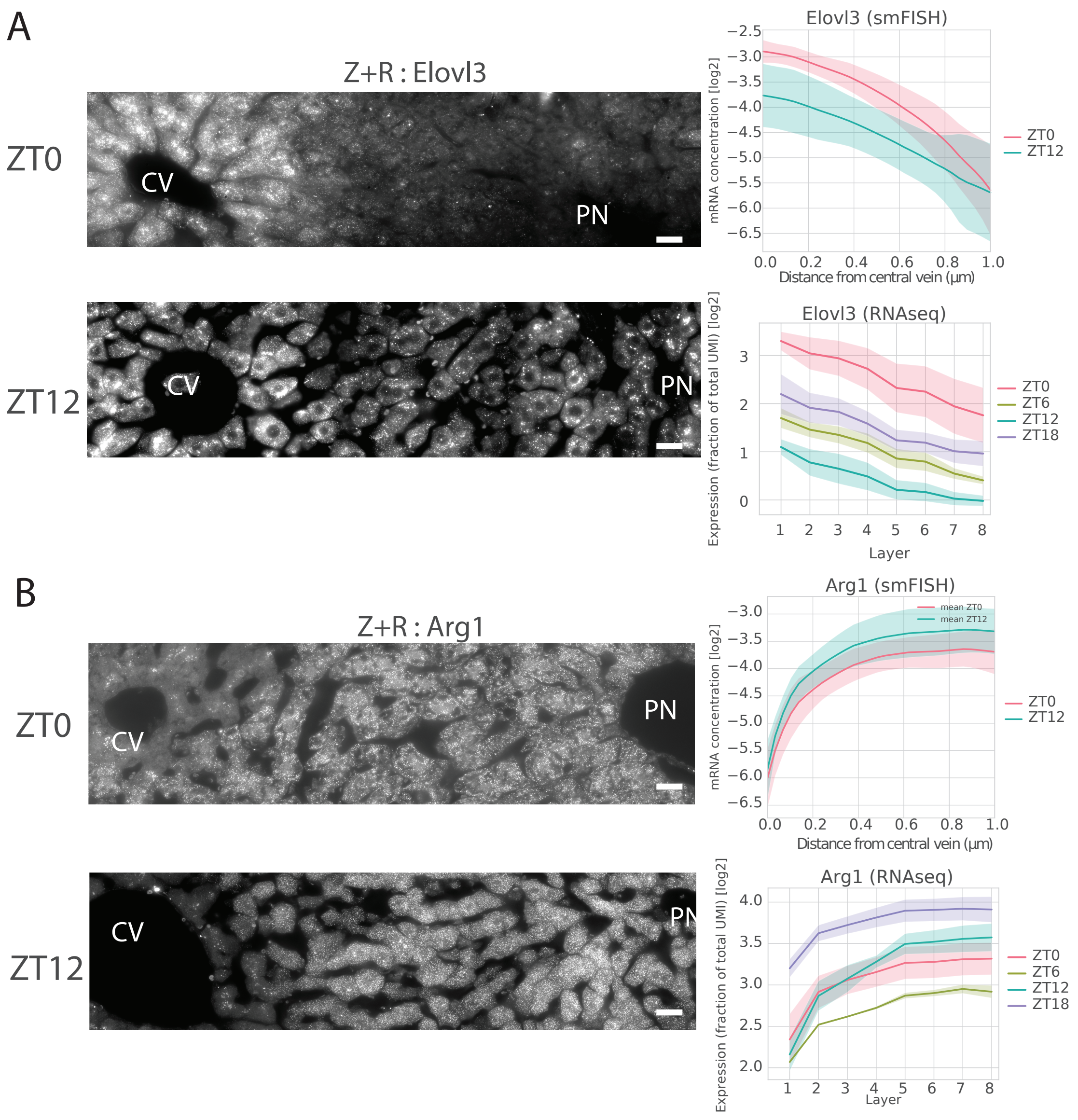


**Supplementary Figure 4: Additional validations for the Z+R category** (**A-B**) smFISH (Stellaris, Methods) for *Elovl3* (Z+R) and *Arg1* (ZxR). smFISH quantifications were made for ZT0 and ZT12 (Methods). **Left**: representative images at ZT0, ZT12 for *Elovl3* (A) or *Arg1* (B). Pericentral veins (CV) and a periportal node (PN) are marked. Scale bar - 20$\mu$m. **Right:** quantified profiles for each gene in the two time points from smFISH (top, shaded area indicate SD across images), and scRNA-seq data (bottom, shaded areas is SD across mice).

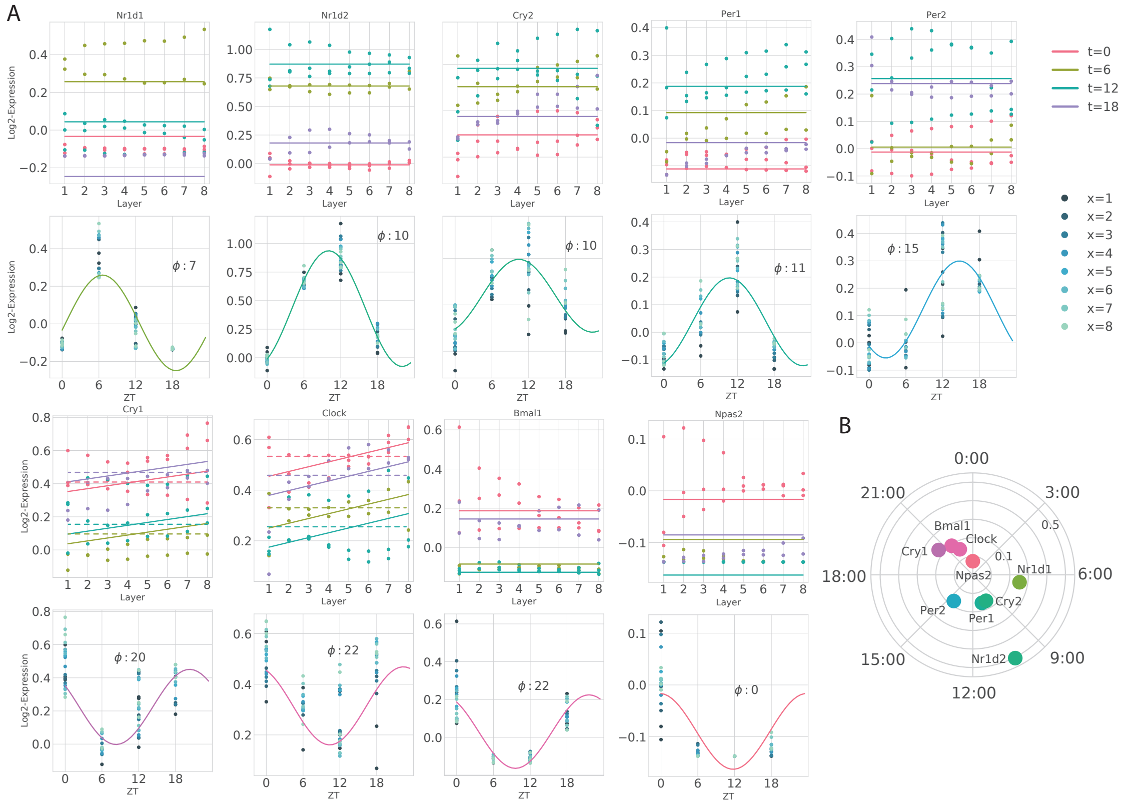


**Supplementary Figure 5: the core circadian-clock is not zonated.** (**A**) Spatial and temporal profiles and fits for circadian core-clock genes. Peak times are indicated on the temporal representation. For the genes *Cry1* and *Clock*, additional dashed lines represent fits for the R model, as the Schwartz BIC weights from the R and Z+R models were close (Supplementary Table S2). (**B**) Amplitudes and peak times of the core-clock circadian genes in a polar coordinate representation (clock-wise ZT times are indicated, distance from the center corresponds to the amplitude) show the expected organization of core clock transcript in the liver.

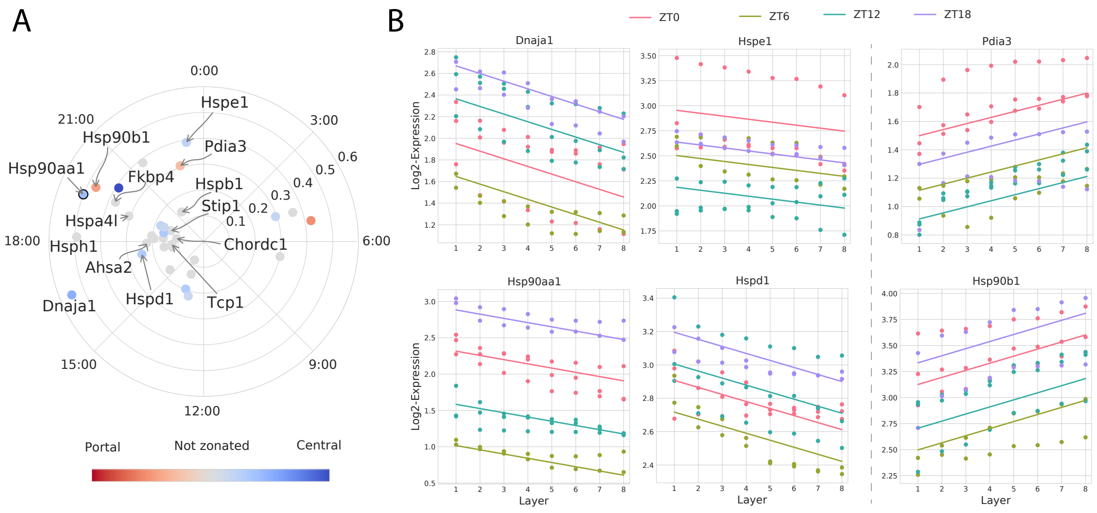


**Supplementary Figure 6: Spatio-temporal mRNA expression profiles for heat-responsive genes. Represented genes correspond to the top 200 targets from the ChIP-Atlas (http://chip-atlas.org/) list for mouse HSF1. (A)** Polar plot representation of the transcripts that are R, Z+R, or ZxR among the HSF1 targets. Genes involved in chaperone functions (chaperones, co-chaperones, or chaperone facilitators) are named. Color indicates zonation, while grey dots show purely rhythmic genes. **(B)** Spatial representation of the transcripts corresponding to proteins involved in chaperone functions, separated in central (left, cytoplasmic function) and portal (right, endoplasmic reticulum function) zonation.

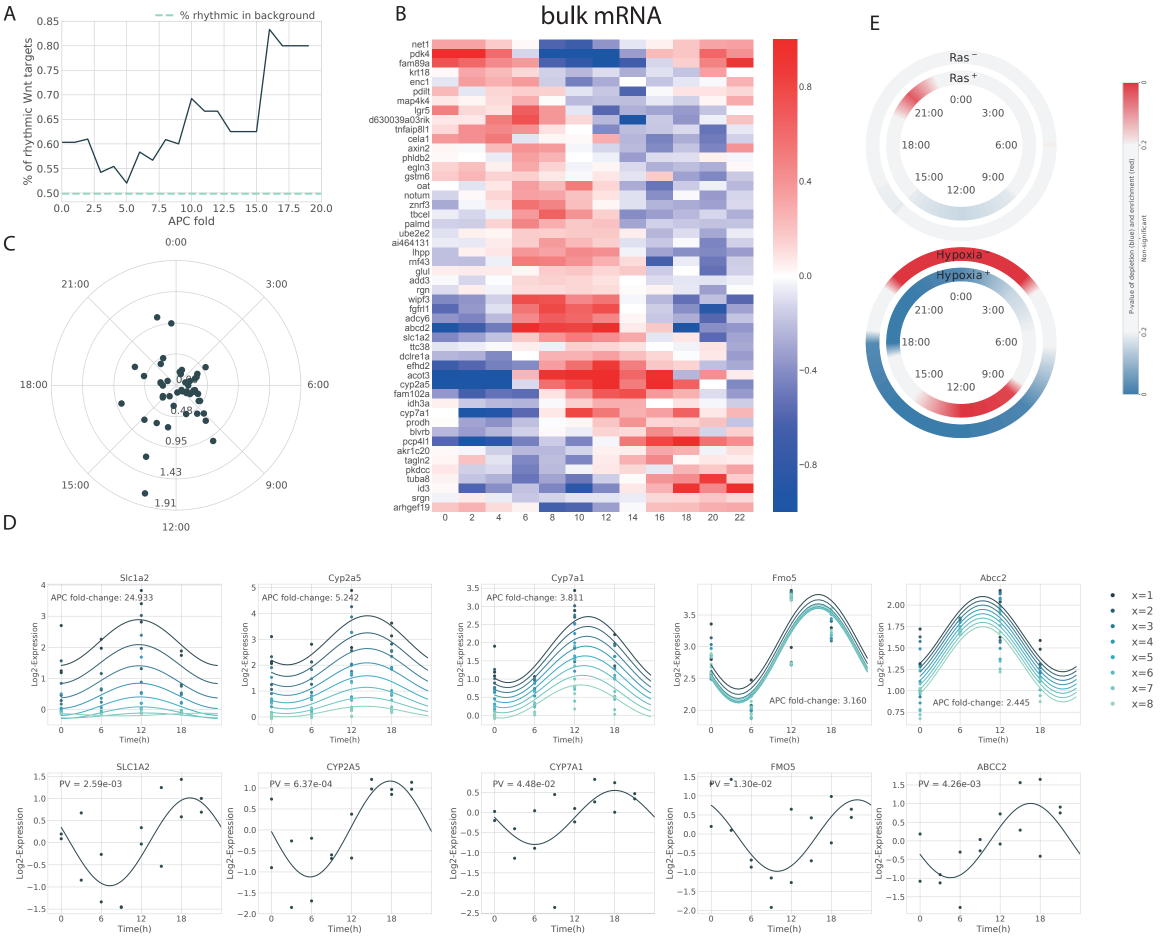


**Supplementary Figure 7: Rhythmicity of Wnt targets in bulk RNA-seq, and proteomics liver time series data (*Robles et al., 2014*).** **(A)** Enrichment of rhythmic genes (R, Z+R and ZxR) among the targets of the Wnt pathway, computed on the bulk dataset (*Atger et al. 2015*). Targets above a given percentile (x-axis) of Apc-KO fold change are considered. The percentage of rhythmic genes in the whole *Atger et al.* dataset is indicated by a dashed blue line. **(B)** Bulk mRNA (coming from *Atger et al.* dataset) rhythmicity profiles of Wnt targets among the top-50 targets with highest Apc-KO fold change. Gene profiles are centered around their mean. An enrichment of the phases around ZT8-14 is observed, in agreement with Figure 6A. **(C)** Polar plot representation of the individual gene phases and amplitudes represented in panel B (bulk data). **(D)** Temporal representation of selected genes profiles from the scRNA-seq (top) and bulk proteomics (bottom, from *Robles et al, 2014*) data. Represented profiles are the ones with (1) the highest Apc-KO fold change, (2) a significantly rhythmic protein (pinf0.05), and (3) belonging to the Z+R or ZxR category. **(E)** Enrichment/depletion at different times (window size: 3h), of both positive and negative Ras and Hypoxia targets (background: all R and Z+R genes). Colormap shows p-values (two-tailed hypergeometric test): red (blue) indicates enrichment (depletion).

#### Part II

### Supplementary Table captions

**Supplementary Table 1. Characteristics of fitted gene profiles.** Model classification for each category (one sheet per category) and each gene (column A, 'Name'). Parameter values of the best model are provided in columns B to H ('mu0' to 'b1'). The standard deviation of the amplitude and phase across the different layers (relevant for ZxR) are provided in columns I and J ('Amplitude spread' and 'Phase spread'). Mean expression profiles in natural and log-transformed scale are provided in columns K and L ('Average expression' and 'Average transformed expression'). The total variance per data point of the profile (computed as the sum of squared distances of the data points to the mean, divided by the number of data points) is provided in column M ('Total Variance'). This number can be compared to the variance per data point explained by the model when keeping only fixed effects (column N, 'Explained variance (fixed effect/data point)', computed as the sum of squared distances of the model predictions to the model mean, when keeping only fixed effects, divided by the number of data points), or both fixed and random effects (column O, 'Explained variance (fixed and random effect/data point)', computed as previously but keeping both fixed and random effects). The relative explained variance (column P, 'Relative Explained variance (fixed effects)', computed as the ratio between column N and M) provides a measure, between 0 and 1, of how well the fixed effects of the model explain the data. Genes are sorted by explained variance.

**Supplementary Table 2. Probability of each model.** Probability of each model, for each gene (column A, 'Name'). The probabilities for the different models are provided in columns B to F. Probabilities are computed as Schwartz BIC weights.

**Supplementary Table 3. KEGG analysis using EnrichR.** KEGG pathway classification, for different combinations of categories (sheets name). Available categories include flat/noisy genes, all zonated genes (Z and Z+R), central (Zc and Zc+R) and portal genes (Zp and Zp+R), central (Zc+R) and portal zonated-rhythymic genes (Zp+R), and purely rythmic (R) genes. KEGG Pathways (column A) having a significant p-value ($< 0.1$) are represented. An ajdusted p-value is provided with each (column B), with the corresponding OR (column C). Finally, the genes observed in the current category are listed in subsequent columns (with explicit headers, column index is sheet-dependent). For zonated genes, the percentage of central genes observed in the category is also provided; while for rhythmic gene, the phase of the genes belonging to the categories are provided.
